# 4’-Fluorouridine is a broad-spectrum orally efficacious antiviral blocking respiratory syncytial virus and SARS-CoV-2 replication

**DOI:** 10.1101/2021.05.19.444875

**Authors:** Julien Sourimant, Carolin M Lieber, Megha Aggarwal, Robert M Cox, Josef D Wolf, Jeong-Joong Yoon, Mart Toots, Chengin Ye, Zachary Sticher, Alexander A Kolykhalov, Luis Martinez-Sobrido, Gregory R Bluemling, Michael G Natchus, George R Painter, Richard K Plemper

**Affiliations:** Institute for Biomedical Sciences, Georgia State University, Atlanta, GA 30303, USA; Texas Biomedical Research Institute, San Antonio, TX 78227, USA; Emory Institute for Drug Development, Emory University, Atlanta, GA 30322, USA; Drug Innovation Ventures at Emory (DRIVE), Atlanta, GA 30322, USA; Department of Pharmacology, Emory University School of Medicine, Atlanta, GA 30322, USA; Department of Pediatrics, Emory University School of Medicine, Atlanta, GA 30322, USA

## Abstract

The COVID-19 pandemic has underscored the critical need for broad-spectrum therapeutics against respiratory viruses. Respiratory syncytial virus (RSV) is a major threat to pediatric patients and the elderly. We describe 4’-fluorouridine (4’-FlU, EIDD-2749), a ribonucleoside analog that inhibits RSV, related RNA viruses, and SARS-CoV-2 with high selectivity index in cells and well-differentiated human airway epithelia. Polymerase inhibition in *in vitro* RdRP assays established for RSV and SARS-CoV-2 revealed transcriptional pauses at positions *i* or *i*+3/4 post-incorporation. Once-daily oral treatment was highly efficacious at 5 mg/kg in RSV-infected mice or 20 mg/kg in ferrets infected with SARS-CoV-2 WA1/2020 or variant-of-concern (VoC) isolate CA/2020, initiated 24 or 12 hours after infection, respectively. These properties define 4’-FlU as a broad-spectrum candidate for the treatment of RSV, SARS-CoV-2 and related RNA virus infections.

**One-Sentence Summary:** 4’-Fluorouridine is an orally available ribonucleoside analog that efficiently treats RSV and SARS-CoV-2 infections *in vivo*.

## Main Text

The COVID-19 experience has demonstrated that proactively developed orally-bioavailable broad-spectrum antivirals are likely the best option to allow rapid response to a newly emerging viral pathogen. Remdesivir, a direct-acting broad-spectrum antiviral, is still the only approved therapeutic for use against SARS-CoV-2 infection, but its dependence on intravenous administration has compromised its clinical impact (*1*). We have demonstrated efficacy of orally available EIDD-2801 (molnupiravir) against influenza viruses in human organoid models and ferrets (*2*), and using the ferret model were subsequently first to show that antiviral efficacy of molnupiravir extends to SARS-CoV-2 *in vivo* (*3*). Initial human data for molnupiravir were encouraging (*4*) and the drug is in advanced clinical trials for treatment of COVID-19. However, even this highly accelerated development timeline reiterates that, to have a substantial impact on a mounting pandemic, an antiviral must be approved for human use before a new pathogen emerges.

We have identified RSV disease as a viable primary indication to develop a broad-spectrum antiviral drug, based on the unmet major health threat imposed by RSV and well-established protocols for clinical trials of novel anti-RSV therapeutics. RSV infections cause over 58,000 hospitalizations of children below five years of age in the United States annually, and approximately 177,000 hospitalizations of adults above the age of 65 (*5–8*). Despite this major health and economic burden, no therapeutics have been licensed specifically for treatment of RSV disease (*9*).

Anti-RSV drug discovery efforts have increasingly focused on inhibiting the viral RNA-dependent RNA polymerase (RdRP) complex (*10*). The core polymerase machinery comprises the large (L) polymerase protein, its obligatory cofactor, the phosphoprotein (P), and the encapsidated negative-sense RNA genome (*10*). Allosteric inhibitors of RSV L have demonstrated potent antiviral activity as seen, for instance, with the experimental drug candidate AVG-233 (*11*) and inhaled PC786 (*12*). However, the antiviral indication spectrum of these inhibitors is narrowly focused on RSV. To improve future pandemic preparedness, a broadened activity spectrum reaching beyond the original licensing indication has emerged as an important objective for the development of next generation antivirals. Narrow therapeutic windows of acute viral diseases (*13*) furthermore require drug administration early after infection, underscoring that next-generation therapeutics against respiratory viruses must be orally bioavailable to meet the time constraints of effective antiviral therapy.

In search of suitable candidates meeting these objectives, we examined 4’-FlU. The development of balapiravir, a prodrug of 4’-azidocytidine, against HCV and other flavivirus infections highlights that 4’-substituted ribonucleosides have antiviral promise (*14, 15*). Fluoro-substitutions appeared furthermore particularly promising, since the small atomic radius and strong stereo-electronic effect of fluorine can influence the ribose backbone conformation, impacting intra- and intermolecular interactions. Having established broad-spectrum antiviral activity of 4’-FlU in *ex vivo* models and determined its mechanism of action (MOA), we validated oral efficacy in two different animal models, the RSV mouse model and the SARS-CoV-2 ferret model. This study introduces 4’-FlU as a promising next-generation antiviral candidate with broad-spectrum potential that may provide a therapeutic option against RSV disease and aid in increasing pandemic preparedness.

## Results

Having recognized the unmet need of RSV disease as a valid primary indication to advance a novel broad-spectrum antiviral, we assessed activity of 4’-FlU (Fig. 1A) on immortalized HEp-2 cells against a recombinant RSV A2-line19F (recRSV A2-L19F) (*16*) and clinical RSV isolates.

**Fig. 1.**
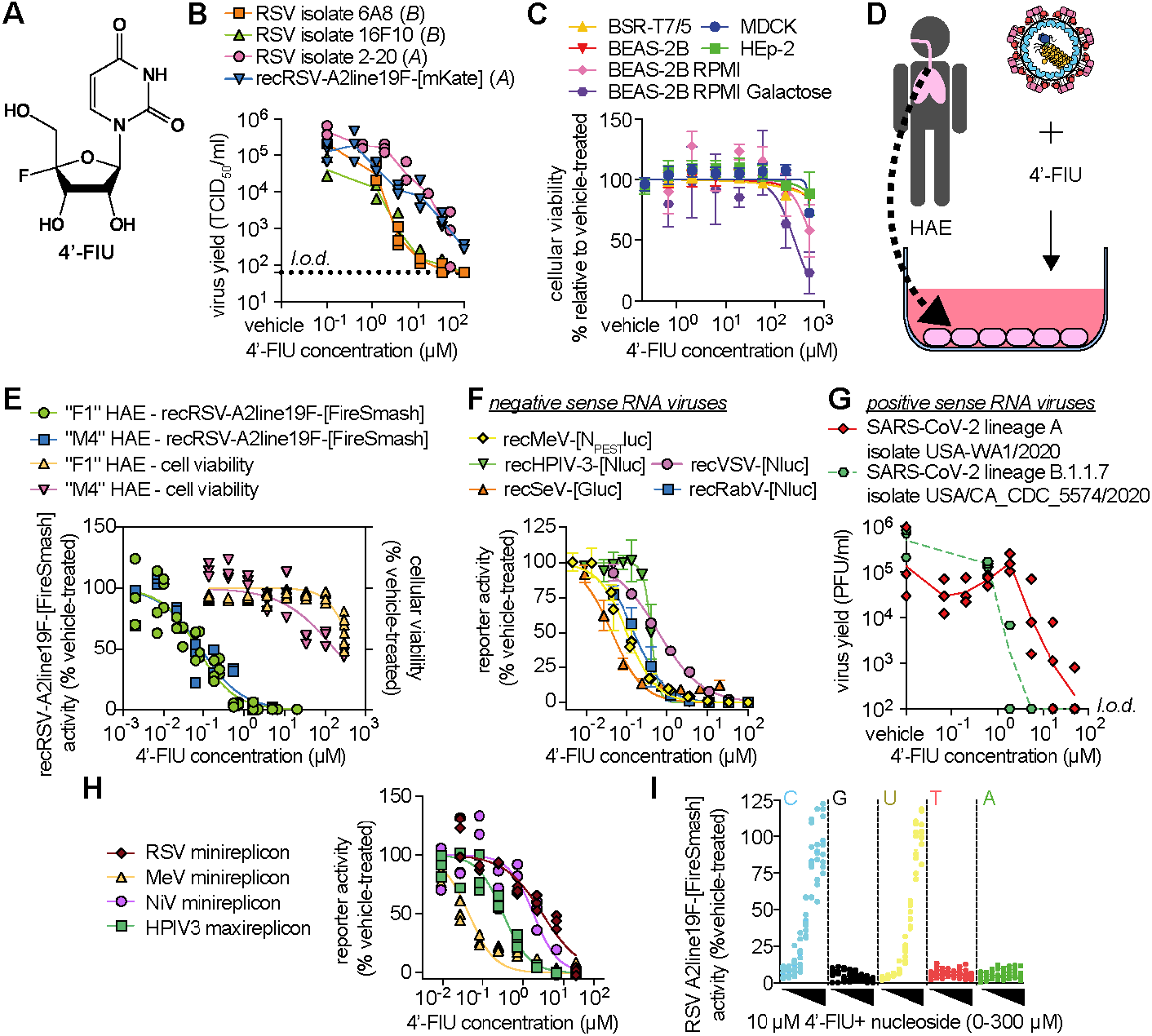
4’-FlU is a potent antiviral with broad spectrum and high selectivity index. **(A)** Chemical structure of 4’-FlU. **(B)** Virus yield reduction of RSV clinical isolates 6A8, 16F10 (B antigenic subgroup), 2-20 and recombinant recRSV-A2line19F-[mKate] (A antigenic subgroup). **(C)** HEp-2, MDCK, BHK-T7 and BEAS-2B cells were assayed for reduction in cell metabolism after incubation with serial dilutions of 4’-FlU or vehicle. (**D,E)** Human airway epithelial (HAE) cells from “F1” and “M4” donors were incubated with serial dilutions of 4’-FlU or vehicle (DMSO) treatment (D), and either infected with recRSV-A2line19F-[FireSMASh] to assess viral inhibition through reporter activity or uninfected to determine cytotoxicity (E). **(F)** Dose-response curves of 4’-FlU against a panel of recombinant mononegaviruses. Viral inhibition determined by reporter activity. **(G)** Virus yield reduction of SARS-CoV-2 WA1/2020 and CA/2020 isolates after incubation with serial dilutions of 4’-FlU or vehicle. **(H)** Dose-response curves of 4’-FlU against transiently expressed viral polymerase complexes from mononegaviruses MeV, RSV, NiV or HPIV-3. **(I)** recRSV-A2line19F-[FireSMASh]-infected cells were treated with 10 μM of 4’-FlU and serial dilutions of exogenous nucleotides in extracellular media. Viral inhibition determined by reporter activity. Symbols represent independent repeats (B,E,G,H,I) or mean with standard deviation (C,F), and lines represent means.

### 4’-FlU is a broad-spectrum mononegavirus inhibitor with high SI

The compound showed potent dose-dependent activity against all RSV strains tested, returning half-maximal efficacious concentrations (EC_50_ values) ranging from 0.61 to 1.2 μM (Fig. 1B) (Table S1). This cell culture potency was on par with anti-RSV activity of previously reported NHC (*17*), the free base of molnupiravir that is currently in clinical development (Fig. S1). Global metabolic activity of established human and animal cell lines (HEp-2, MDCK, BHK-T7, BEAS-2B) exposed to up to 500 μM of 4’-FlU remained unaltered (Fig. 1C) (Table S2). When glucose was replaced with galactose as carbohydrate source to link cell metabolic activity strictly to mitochondrial oxidation (*18*), we determined a half-maximal cytotoxic concentration (CC_50_) of 4’-FlU of 250 μM (Fig. 1C) (Table S2).

When tested on disease-relevant primary human airway epithelial cells (HAE) derived from two different donors (Fig. 1D), 4’-FlU showed ≥17-fold increased anti-RSV potency but unchanged low cytotoxicity (CC_50_ 169 μM) (Fig. 1E), resulting in a high selectivity index (SI = EC_50_/CC_50_) of ≥1877. Consistent with these findings, in-cell quantitative immunocytochemistry confirmed that 4’-FlU similarly reduced steady-state levels of nuclear-(SDH-A; IC_50_ 272.8 μM) and mitochondrial-(COX-I; IC_50_ 146.8 μM) encoded proteins in HAEs only at high concentrations (Fig. S2).

Assessment of the broader 4’-FlU indication spectrum against a panel of related major pathogens of the mononegavirus order, including measles virus (MeV), human parainfluenza virus type 3 (HPIV3), Sendai virus (SeV), vesicular stomatitis virus (VSV), and rabies virus (RabV) consistently demonstrated sub-micromolar active concentrations (Fig. 1F) (Table S1). The positive-sense RNA betacoronavirus SARS-CoV-2 was apparently only slightly less sensitive to 4’-FlU, with EC_50_ values ranging from 0.5 to 5.1 μM against isolates of different lineages (Fig. 1G) (Table S1).

An initial mechanistic characterization of 4’-FlU in cell-based minireplicon systems revealed inhibition of pneumovirus and paramyxovirus RdRP complex activity (Fig. 1H) (Table S1). In accordance with broad-spectrum activity against mononegavirus, the RdRP activity of Nipah virus (NiV), a highly pathogenic zoonotic paramyxovirus with strong pandemic potential (*19*), was also efficiently inhibited by 4’-FlU in an NiV minireplicon reporter assay. The antiviral effect of 4’-FlU was dose-dependently reversed by addition of an excess of exogenous pyrimidines, cytidine and uridine, but not purines, to the cultured cells, which furthermore is consistent with competitive inhibition of RdRP activity (*2*) (Fig. 1I).

### Incorporation of 4’-FlU by RSV and SARS-CoV-2 RdRP causes a sequence-modulated transcriptional pause

To characterize the molecular MOA of 4’-FlU, we purified recombinant RSV L and P proteins expressed in insect cells (Fig. 2A) and characterized the performance of the bioactive 5’-triphosphate form of 4’-FlU (4’-FlU-TP) in *in vitro* primer extension assays (*20*) (Fig. 2B). In the presence of radio-labelled ATP and an increasing amount of UTP, purified RSV RdRP complexes elongated the primer to the third position, which called for CTP, and continued further upon addition of CTP (Fig. 2C) (Fig. S3) (Data S1). Replacing UTP with 4’-FlU-TP likewise resulted in efficient primer extension up to the third nucleotide, confirming that RSV RdRP recognizes and incorporates 4’-FlU as uridine nucleotide (Fig. 2C). Incorporation kinetics (*21*) showed a moderate reduction of substrate affinity for 4’-FlU-TP compared to UTP, reflected by Km values of 24.9 μM versus 6.71 μM (Fig. 2D). These findings confirmed efficient incorporation of 4’-FlU-TP by RSV RdRP. Further addition of CTP to the reaction mix resulted in limited elongation to gacgcAA(4’-FlU)CA(4’-FlU)A rather than the expected full-length gacgcAA(4’-FlU)CA(4’-FlU)AACA, which suggested delayed chain termination by incorporated 4’-FlU (Fig. 2C) (Data S1).

**Fig. 2.**
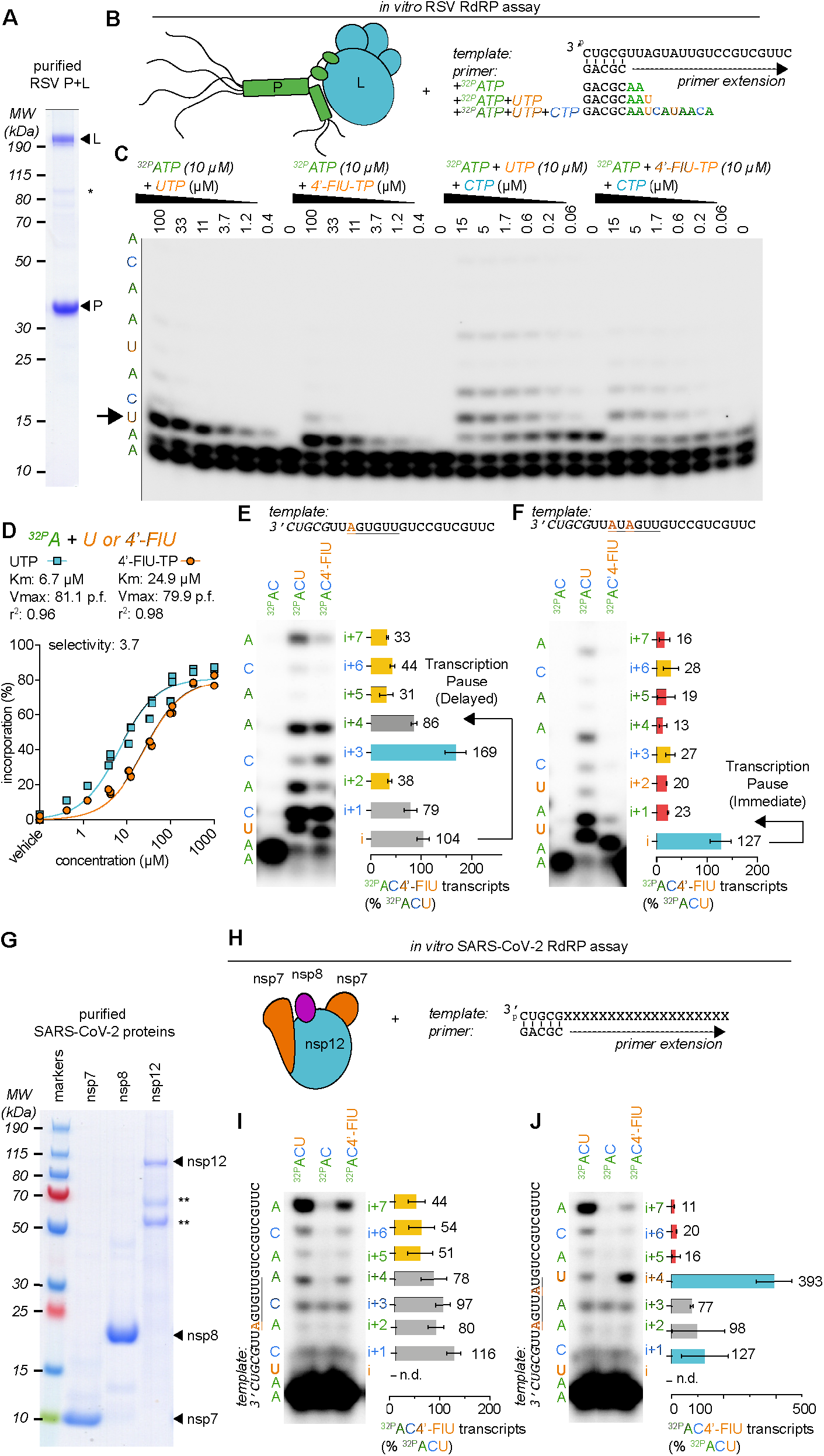
4’-FlU induces a delayed stalling of phosphodiester bond formation by RSV and SARS-CoV-2 RdRP. **(A)** SDS-PAGE with Coomassie blue staining of purified recombinant RSV RdRP complexes (L and P proteins) used for *in vitro* RdRP assays. Stars denote cellular contaminants. **(B)** Schematics of the primer extension assay. **(C)** Urea-PAGE fractionation with single nucleotide resolution of RNA transcripts produced through primer extension by the RSV RdRP in the presence of the indicated nucleotides. Arrow highlights the site of first incorporation of 4’-FlUTP (n=3). **(D)** Kinetic analysis of autoradiographs from (C). Non-linear regression with Michaelis-Menten model. Km and Vmax with 95% confidence intervals (CIs) and goodness of fit (r^2^) are indicated (n=3). (**E,F)** Urea-PAGE fractionation of RNA transcripts produced by RSV RdRP in the presence of the indicated templates and nucleotides. 4’-FlU-TP bands were normalized to the corresponding band after UTP incorporation. Bars represent mean and error bar represent standard deviation (n=3). (**G)** Purified recombinant SARS-CoV-2 RdRP complexes (nsp7, 8, and 12 proteins) used for *in vitro* RdRP assays as in (A). (**H-J)** Urea-PAGE fractionation of RNA transcripts produced by SARS-CoV-2 RdRP in the presence of the indicated templates and nucleotides.

When a modified template called for incorporation of only a single UTP (Fig. 2E)(Data S1), we noted accumulation of signal at position *i*+3 after 4’-FlU-TP, whereas the efficiency of full primer elongation (*i*+7) was strongly reduced compared to extension after UTP. However, repositioning the incorporation site further downstream in the template triggered polymerase pause at position *i* (Fig. S4), suggesting template sequence dependence of the inhibitory effect. Transcription pauses at *i* or *i*+3 were also observed after multiple 4’-FlU incorporations: a UxUxxx template (Fig. 2F) and direct tandem incorporations (Fig. S4) caused termination at position *i*, whereas increasing spacer length between the uridines to two, three, or four nucleotides resulted in increasingly pronounced pausing at *i*+3 (Fig. S4). This sequence-dependent, variable delayed polymerase termination within one to four nucleotides of the incorporation site was equally prominent when we examined *de novo* initiation of RNA synthesis at the promoter using a synthetic native RSV promoter sequence rather than extension of primer-template pairs (Fig. S5).

Purification of a core SARS-CoV-2 polymerase complex (non-structural proteins (nsp) 7, 8 and 12) from bacterial cell lysates (Fig. 2G) and assessment of RdRP bioactivity in equivalent primer-extension *in vitro* polymerase assays (Fig. 2H) demonstrated incorporation of 4’-FlU-TP in place of UTP by the coronavirus RdRP (Fig. 2I), again with no sign of direct chain termination. While transcriptional pauses of SARS-CoV-2 RdRP were less noticeable than with RSV RdRP upon incorporation of 4’-FlU, polymerase pausing and/or termination was again triggered after multiple incorporations of 4’-FlU-TP in a sequence-dependent manner, and were particularly prominent when subsequent incorporation of 4’-FlU-TP occurred at *i*+4 position (Fig. 2J) (Fig. S6). No primer extension occurred when the nsp12 subunit was omitted or an nsp12 variant carrying mutations in the catalytic site was used, confirming specificity of the reaction (Fig. S6) (Data S1).

### 4’-FlU is rapidly anabolized, metabolically stable, and potently antiviral in disease-relevant well-differentiated HAE cultures

Quantitation of 4’-FlU and its anabolites in primary HAE cells after exposure to 4’-FlU for different times (Fig. 3A) demonstrated rapid intracellular accumulation of 4’-FlU, reaching a level of 3.42 nmol/million cells in the first hour of exposure (Fig. 3B). Anabolism to bioactive 4’-FlU-TP was likewise efficient, resulting in concentrations of 10.38 nmol/million cells at peak (4 hours after exposure start) and 1.31 nmol/million cells after 24 hours at plateau. Subsequent wash-out of the compound revealed high metabolic stability of the anabolite, which remained present in sustained concentrations of approximately 1 nmol/million cells over a 6-hour monitoring period, corresponding to an extrapolated half-life of 9.7 hours (Fig. 3C).

**Fig. 3.**
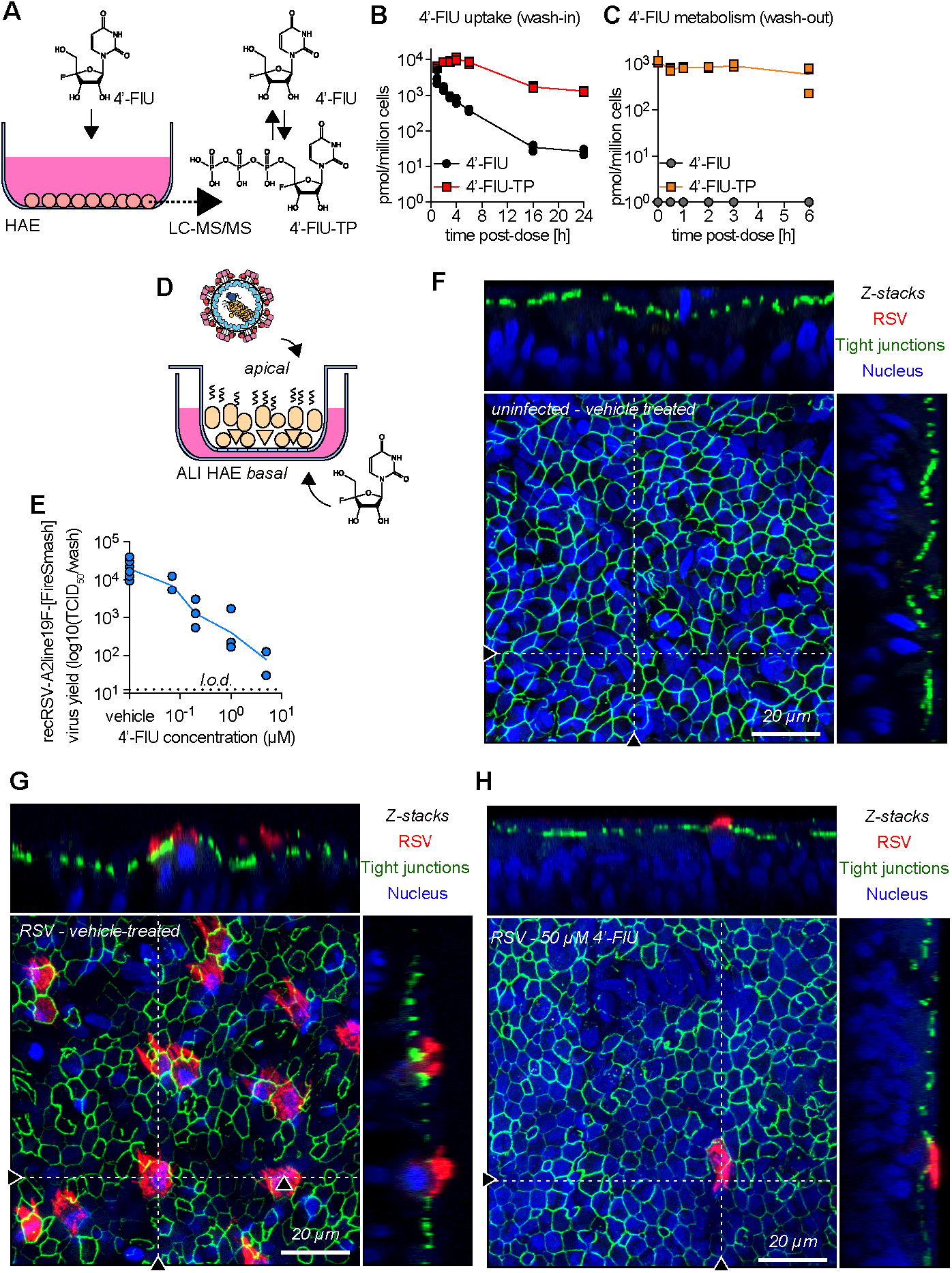
4’-FlU is efficiently metabolized to 4’-FlU-TP in HAE cells and demonstrate potent efficacy in a human airway surrogate model. **(A-C)** Cellular uptake and metabolism of 4’-FlU in HAE cells quantified by mass spectrometry (A). Intracellular concentration of tri-phosphorylated 4’-FlU after either exposure to 20 μM 4’-FlU for 0,1, 2, 3, 4, 6,16, and 24 hours (B), or a first 24-hour incubation followed by removal of compound for 0, 0.5, 1, 2, 3, 6 hours before quantification (n=3). **(D)** HAE cells were cultivated at air-liquid interface (ALI) to serve as a surrogate model of human lung tissues exposed to respiratory pathogens on the apical side and to treatment on basal side. **(E)** Virus yield reduction of recRSV-A2line19F-[FireSMASh] shed from the apical side in ALI HAE after incubation with serial dilutions of 4’-FlU on the basal side (n=3). **(F-H)** Confocal microscopy of ALI HAE cells infected with recRSV-A2line19F-[FireSMASh], at 5 days post-infection. RSV infected cells, tight junctions and nuclei were stained with anti-RSV immunostaining, anti-ZO-1 immunostaining and Hoechst 34580. z-stacks of 30 μm slices (1 μm thick) with 63× objective with oil immersion. Dotted lines represent the location of x-z and y-z stacks; scale bar: 20 μm. In all panels, symbols represent independent biological repeats and lines represent means.

To explore efficacy in a disease-relevant human tissue model, we cultured the HAEs at air-liquid interface, inducing the formation of a well-differentiated 3D epithelium that mimicked hallmarks of a human airway epithelium, including ciliated and mucus-producing cells (*22*) (Fig. 3D). When added to the basolateral chamber of the transwells after apical infection of the epithelium with RSV, 4’-FlU potently reduced apically shed progeny virus titers with an EC_50_ of 55 nM (Fig. 3E). Overall titer reduction spanned nearly four orders of magnitude, ranging from 3.86×10^4^ TCID_50_ in the presence of vehicle volume equivalents to 78.18 TCID_50_ at 5 μM basolateral 4’-FlU, which approached the level of detection.

Confocal microscopy validated formation of a pseudostratified organization of the epithelium with tight junctions in the airway epithelium tissue model (Fig. 3F), visualized efficient RSV replication in vehicle-treated tissue models (Fig. 3G), and confirmed near-sterilizing antiviral efficacy in the presence of 50 μM basolateral 4’-FlU (Fig. 3H) (Fig. S7, S8). Under these conditions, positive staining for RSV antigen was rarely detected.

### 4’-FlU is orally efficacious in a therapeutic dosing regimen in a small-animal model of RSV infection

To test 4’-FlU efficacy *in vivo*, we employed the mouse model of RSV infection, challenging animals with recRSV-A2-L19F, which efficiently replicates in mice (*16*). In a dose-to-failure study, we infected BALB/cJ mice intranasally and initiated once-daily oral treatment two hours after infection at 0.2, 1, or 5 mg 4’-FlU/kg body weight. Treatment at all dose levels resulted in a statistically significant reduction in lung virus load compared to vehicle-treated animals (Fig. 4A). The antiviral effect was dose-dependent and approached nearly two orders of magnitude at the 5 mg/kg dose level, the greatest reduction that we have ever observed in this model (*17*). Consistent with the observed high metabolic stability in HAEs, a twice-daily dosing regimen did not significantly enhance efficacy (Fig. S9). Since animal appearance, body weight, temperature (Fig. S10), and relative lymphocyte and platelet counts (Fig. 4B) (Fig. S11) were unchanged in the 5 mg/kg group compared to vehicle-treated animals, we selected this dose level for the following studies.

**Fig. 4.**
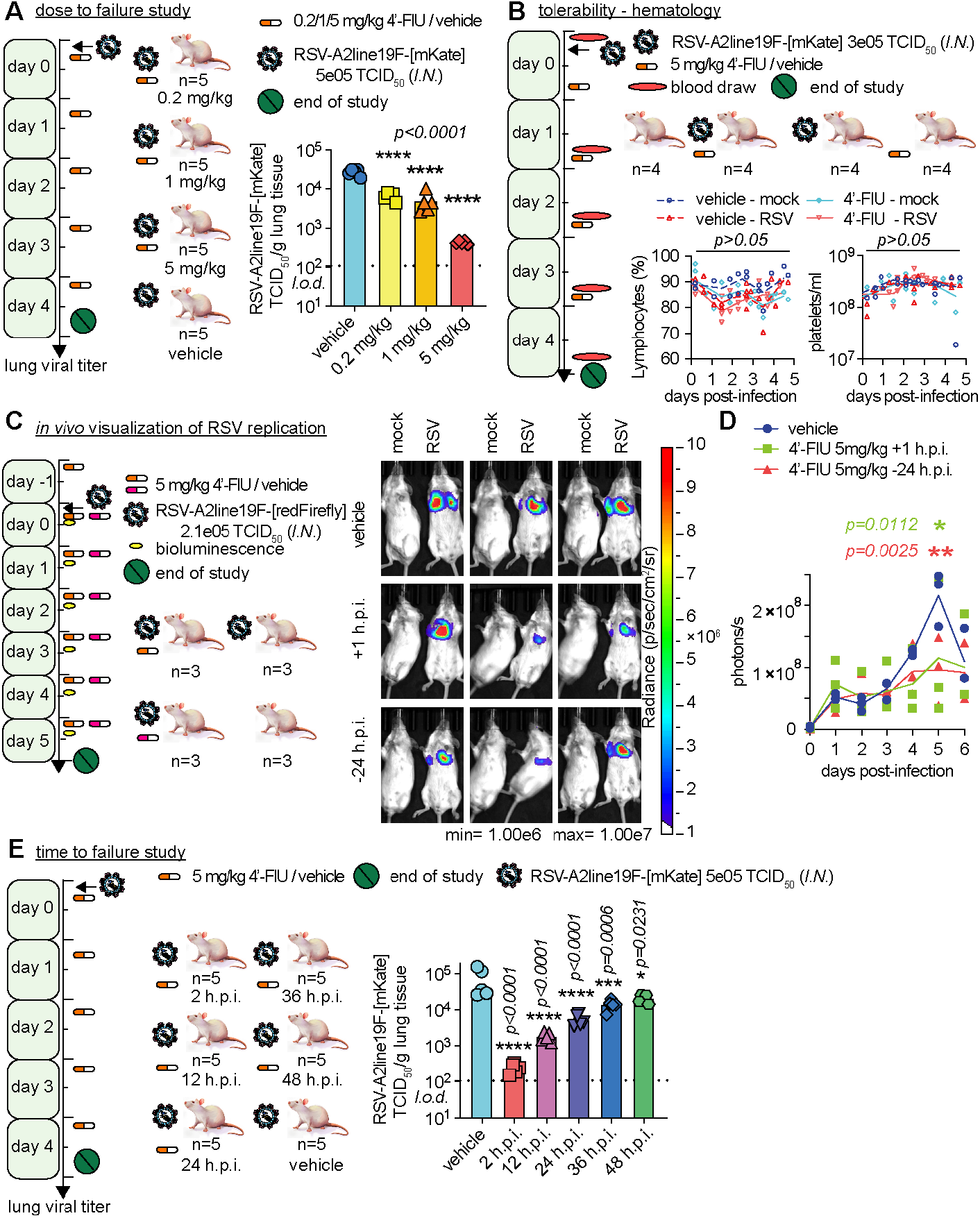
Therapeutic oral efficacy of 4’-FlU against RSV replication in mice. **(A)** Balb/cJ mice were inoculated with recRSV-A2line19F-[mKate] and treated as indicated. At 4.5 days after infection, viral lung titers were determined with TCID_50_ titration (n=5). **(B)** Balb/cJ mice were inoculated with recRSV-A2line19F-[mKate] or mock-infected, and treated as indicated. Blood samples were collected prior to infection and at 1.5, 2.5, 3.5, and 4.5 days after infection and lymphocytes proportions with platelets/ml are represented over time (n=4). **(C)** Balb/cJ mice were inoculated with recRSV-A2line19F-[redFirefly] and treated as indicated. *In vivo* luciferase activity was measured daily. **(D)** Total photon flux from mice lungs from (C) over time (n=3). **(E)** Balb/cJ mice were inoculated with recRSV-A2line19F-[mKate] and treated as indicated. At 4.5 days after infection, viral lung titers were determined with TCID_50_ titration (n=5). In all panels, symbols represent individual values and bar or lines represent means. One-way ordinary ANOVA with Tukey’s post-hoc multiple comparisons (B,I) or two-way ANOVA with Dunnett’s post-hoc multiple comparison (C,G). h.p.i.= hours post-infection.

For a longitudinal assessment of therapeutic benefit in individual animals, we employed an *in vivo* imaging system (IVIS) using a red-shifted luciferase-expressing RSV reporter virus (*23*) that we had generated for this study to maximize signal intensity. Daily imaging (Fig. 4C) (Fig. S12) revealed significant reduction of bioluminescence intensity in lungs of 4’-FlU-treated animals, independent of whether treatment was initiated 24 hours prior to, or 1 hour after, infection (Fig. 4D). This IVIS profile is consistent with reduced viral replication and ameliorated viral pneumonia in the treated animals.

To probe the therapeutic window of 4’-FlU, we initiated treatment at 2, 12, 24, 36, and 48 hours after infection. All treatment groups showed a statistically significant reduction of lung virus burden compared to vehicle-treated animals, but effect size was dependent on the time of treatment initiation (Fig. 4E) (Fig. S13). Based on our experience with therapeutic intervention with related respiratory RNA viruses that cause lethal disease (*22*), we consider a reduction of lung virus load of at least one order of magnitude required for robust therapeutic benefit. With this request, the therapeutic window of 4’-FlU extended to 24 hours after infection in mice.

### 4’-FlU is orally efficacious against SARS-CoV2 infection in the ferret model

We next tested efficacy against SARS-CoV-2 WA1/2020 in the HAE organoid models grown at air-liquid interface. Independent of the human donor, SARS-CoV-2 replicated efficiently in these disease-relevant tissue models (Fig. 5A-C)(Fig. S14). Treatment of the infected cultures with basolateral 4’-FlU dose-dependently reduced apical virus shedding, albeit approximately 45-fold less potently (EC_50_: 2.47 μM) than inhibition of RSV and with a limited maximal effect size of approximately two orders of magnitude at 50 μM (Fig. 5D). Confocal microscopy revealed that most of the epithelium was negative for staining of the SARS-CoV-2 nucleocapsid protein under these conditions (Fig. 5E), with only sporadic exceptions of a small number of remaining infected ciliated cells (Fig. 5F)(Fig. S14).

**Fig. 5.**
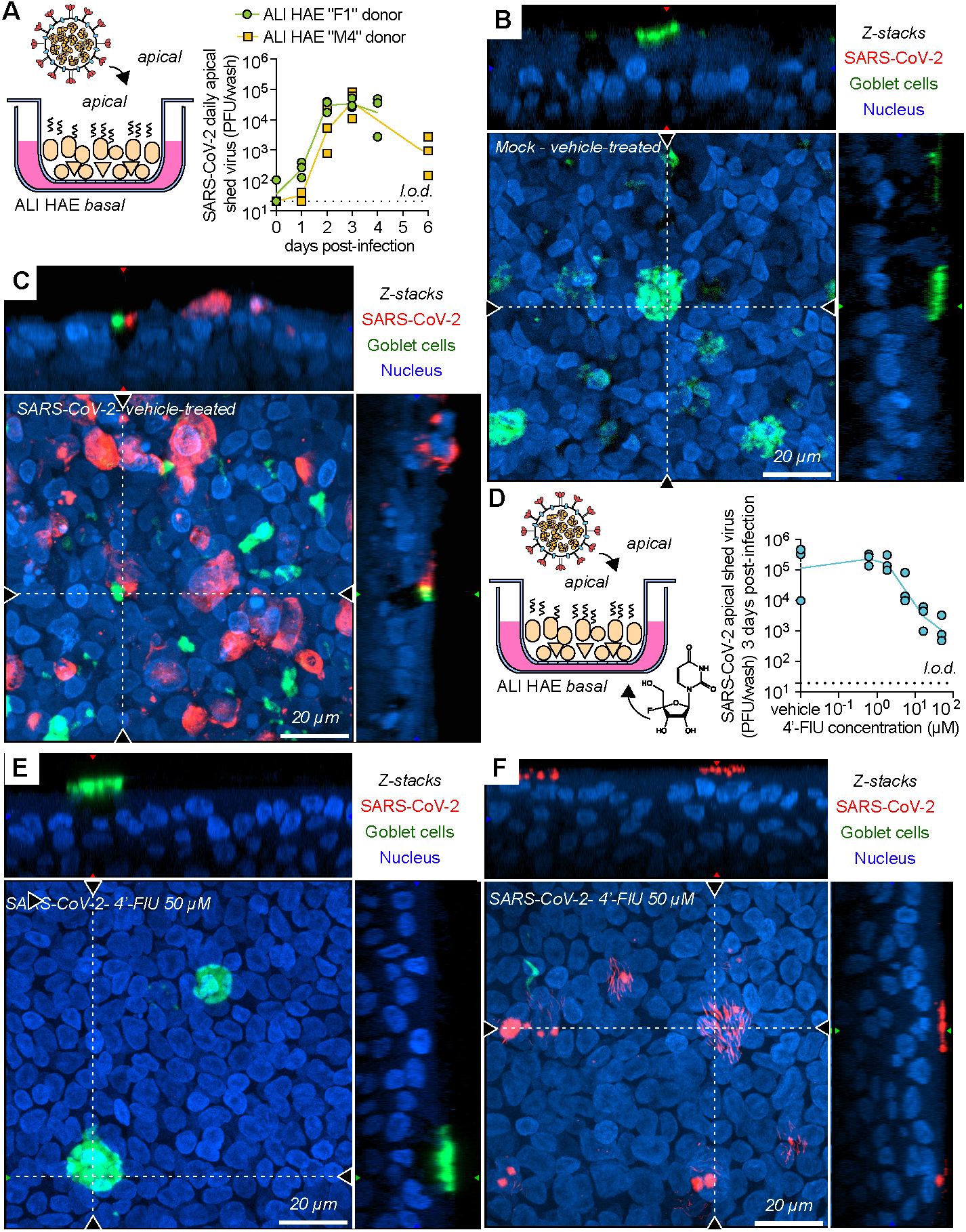
Efficacy of 4’-FlU against SARS-CoV-2 replication in ALI. **(A)** Multicycle growth curve of SARS-CoV-2 WA1/2020 isolate on ALI HAE from two donors. Shed virus was harvested daily and tittered by plaque assay (n=3). **(B,C)** Confocal microscopy of ALI HAE cells mock-infected (B) or infected (C) with SARS-CoV-2 WA1/2020 isolate, at 3 days post-infection. SARS-CoV-2 infected cells, goblet cells and nuclei were stained with anti-SARS-CoV-2 N immunostaining, anti-MUC5AC immunostaining and Hoechst 34580, pseudo-colored in red, green and blue, respectively. z-stacks of 35 μm slices (1 μm thick) with 63× objective with oil immersion. Dotted lines represent the location of x-z and y-z stacks; scale bar: 20 μm. In all panels, symbols represent independent biological repeats and lines represent means. **(D)** Virus yield reduction of SARS-CoV-2 WA1/2020 clinical isolate shed from the apical side in ALI HAE after incubation with serial dilutions of 4’-FlU on the basal side (n=3). **(E,F)** Confocal microscopy of ALI HAE cells infected with SARS-CoV-2 WA1/2020 isolate, and treated with 50 μM 4’-FlU at 3 days post-infection. Rare ciliated cells positive for N are represented in (F).

To assess whether this reduction in viral shedding translates to a significant benefit *in vivo*, we determined efficacy of oral 4’-FlU against 2019-nCoV/USA-WA1/2020 (WA1/2020, A lineage) and the VoC isolate 2019-nCoV/USA/CA_CDC_5574/2020 (CA/2020, B.1.1.7 lineage) in the ferret model, which recapitulates hallmarks of uncomplicated human infection (*3*). For informed dose level selection in ferrets, we first determined single oral dose ferret pharmacokinetic (PK) profiles of 4’-FlU. When administered at 15 or 50 mg/kg, peak plasma concentrations (C_max_) of 4’-FlU reached 34.8 and 63.3 μM, respectively, and overall exposure was 154 ± 27.6 and 413.1 ± 78.1 h×nmol/ml, respectively, revealing good oral dose-proportionality (Fig. 6A) (Table S3). Based on this PK performance, we selected once-daily dosing at 20 mg/kg body weight for efficacy tests (Fig. 6B).

**Fig. 6.**
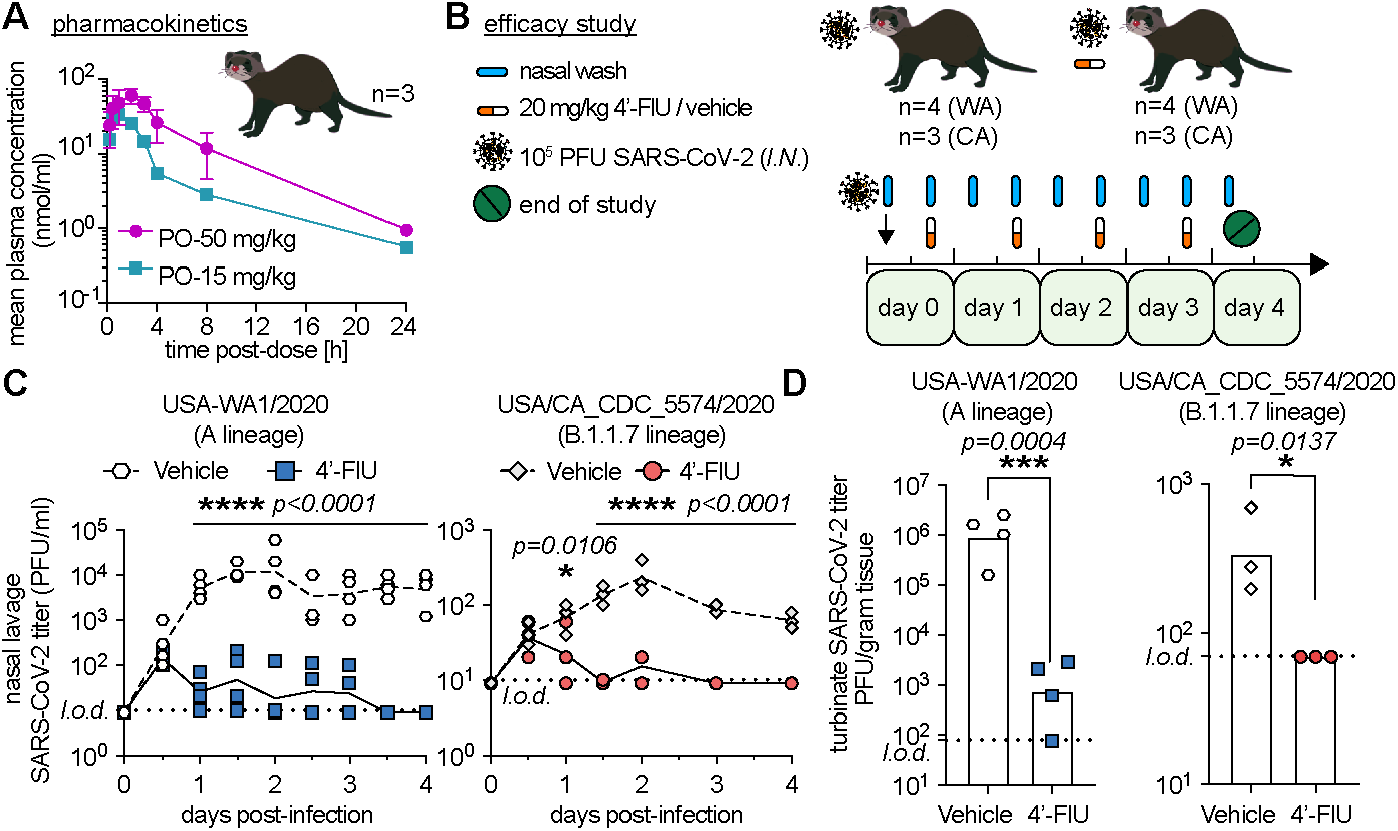
Therapeutic oral efficacy of 4’-FlU against different SARS-CoV-2 isolates in ferrets. **(A)** Single oral dose (15 or 50 mg/kg bodyweight) pharmacokinetics properties of 4’-FlU in ferret plasma (n=3). (**B)** Ferrets were inoculated with SARS-CoV-2 WA1/2020 or CA/2020 isolate and treated as indicated. (**C**) Nasal lavages were performed twice daily and viral titers determined by plaque assay (n=4 (WA1/2020) or n=3 (CA/2020)). (**D)** Viral titers in nasal turbinates at day 4 post-infection. In all panels, symbols represent individual independent biological repeats, lines show mean values. Two-way ANOVA with Dunnett’s post-hoc multiple comparison (C) and unpaired t-test (D).

Intranasal infection of ferrets with 1×10^5^ PFU of either WA1/2020 or CA/2020 resulted in rapid viral shedding into the upper respiratory tract. An upper plateau of shed virus was reached in vehicle-treated animals 48 hours after infection at 1.2×10^4^ PFU/ml and 2.3×10^2^ PFU/ml nasal lavage, respectively (Fig. 6C). Therapeutic treatment with 4’-FlU initiated 12 hours after infection caused a significant reduction in shed virus burden by approximately three orders of magnitude (WA1/2020) to less than 50 PFU/ml nasal lavage and one order of magnitude (CA/2020), respectively, within 12 hours of treatment onset. Virus loads remained low for the remainder of the study. Viral titers in nasal turbinate tissue extracted four days after infection (Fig. 6D) and associated viral RNA copy numbers (Fig. S15) correlated with this reduction in shed virus load. Shedding of infectious particles ceased completely in all animals after 2.5 days of treatment (three days post-infection). These results demonstrate that once-daily oral administration of 4’-FlU efficiently suppresses replication of the original WA1/2020 SARS-CoV-2 isolate and the more recently emerged CA/2020 B.1.1.7 lineage in a relevant animal model.

## Discussion

Our anti-RSV efforts have yielded 4’-FlU, an orally efficacious ribonucleoside analog inhibitor with broad-spectrum activity against negative sense RNA viruses of the pneumovirus, rhabdovirus, and paramyxovirus families, which includes the highly pathogenic henipaviruses that carry high pandemic threat potential (*19*). *In vitro* RdRP assays identified polymerase pausing or delayed chain-termination as the primary MOA of 4’-FlU, which is reminiscent of the antiviral effect of remdesivir (*24, 25*). In contrast to remdesivir, however, 4’-FlU also induced an immediate RdRP stall after incorporation in a sequence-dependent manner. These properties suggest steric hindrance of the RdRP to advance after incorporation of the compound or to accommodate the next incoming nucleotide as the underlying antiviral mechanism of 4’-FlU.

Albeit with reduced potency, activity of 4’-FlU extended to the positive-sense RNA betacoronavirus SARS-CoV-2. The somewhat lower sensitivity of SARS-CoV-2 to the compound compared to RSV could be due to the exonuclease activity of the coronavirus polymerase complex, which reportedly can eliminate ribonucleoside analogs (*26, 27*). However, SARS-CoV-2 polymerase also showed a higher tendency to advance further along the template after 4’-FlU-TP incorporation than RSV RdRP in the *in vitro* RdRP assays, which do not contain the exonuclease functionality. Accordingly, coronavirus RdRP may also have a somewhat greater capacity to tolerate incorporated 4’-FlU and continue polymerization than mononegavirus polymerase complexes.

Since no small-animal model of RSV infection fully recapitulates all aspects of human RSV disease (*28*), we employed two complementary infection models to assess antiviral efficacy of 4’-FlU, *ex vivo* HAE cultures and the *in vivo* mouse model. Results revealed consistent antiviral efficacy in both systems, indirectly confirming also efficient anabolism in disease-relevant primary human tissues, oral bioavailability of the compound in mice, and accumulation of bioactive 4’-FlU-TP at efficacious levels in mouse lung. Ferret PK profiles validated that 4’-FlU exposure levels are high and dose-proportional after oral administration. Although RSV is known to advance rapidly to viral pneumonia in mice, therapeutic treatment with 4’-FlU demonstrated a 24-hour therapeutic window after infection. Since RSV host invasion is slower in humans (*29*), these data support that a viable therapeutic window should exist for efficacious treatment of human RSV infection.

When comparing activity of 4’-FlU against SARS-CoV-2 and RSV in cultured cells and well-differentiated HAEs, anti-SARS-CoV-2 potency was overproportionally lower in the organoid models. RSV tropism in the organoids is restricted to ciliated cells (*30, 31*), whereas SARS-CoV-2 can reportedly be found in both ciliated and goblet cells (*32–37*). Conceivably, 4’-FlU anabolism efficiency could be higher in ciliated than in goblet cells, affecting anti-SARS-CoV-2 impact in this system. Importantly, however, once-daily oral treatment of ferrets infected with WA1/2020 or the recently emerged VoC CA/2020 reduced virus burden to an almost undetectable level within 12 hours of treatment initiation. These data confirm broad-spectrum anti-coronavirus efficacy of 4’-FlU in a relevant animal model and suggest that 4’-FlU has greater potential to remain active against future VoCs that may be less responsive to spike-targeting vaccines or antibody therapeutics.

Formal tolerability studies with 4’-FlU are still pending, but the compound was well tolerated by the human organoid models and efficacious in murids and mustelids. Blood analysis of treated mice revealed no global changes in blood cell counts, indicating that treatment had no negative effect on the hematopoietic system. These results establish 4’-FlU as a much-needed novel broad-spectrum antiviral with confirmed efficacy against major RNA virus pathogens, making it a promising therapeutic option for severe RSV disease and an important contributor to further improving pandemic preparedness.

## Acknowledgments

We thank C. F. Basler for providing Calu-3 cells, D. Waugh for plasmid pRK792 encoding TEV protease (Addgene plasmid #8830), the Georgia State University High Containment Core and the Department for Animal Research for support, and A. L. Hammond for critical reading of the manuscript.

## Author contributions

Conceptualization: MGN, GRP, RKP.

Investigation: JS, RMC, MT, JY, CML, MA, JDW, RKP

Resources: GRB, AAK, LMS, RKP

Visualization: GRB, JS

Validation: JS, RKP

Funding acquisition: RKP

Project administration: MGN, RKP

Supervision: GRP, RKP

Writing – original draft: JS, RKP

Writing – review & editing: JS, RKP

## Data and materials availability

All data are available in the main text or the supplementary materials. Transfer of EIDD-2749 material to other institutions for research purposes is covered by MTAs from Emory University.

## Supplementary Materials

Materials and Methods

Figs. S1 to S17

Tables S1 to S3

Data S1 to S2

