## Supplemental information for "4’-Fluorouridine is a broad-spectrum orally efficacious antiviral blocking respiratory syncytial virus and SARS-CoV-2 replication"

**This PDF file includes:**

Materials and Methods
Figs. S1 to S17
Tables S1 to S3
Captions for Data S1
Captions for Data S2

**Other Supplementary Materials for this manuscript include the following:**

Data S1
Data S2

### Materials and Methods

#### Study design

The objectives of this study were to explore the mechanism of action and preclinical efficacy of the ribonucleoside analog 4'-FlU against RSV and SARS-CoV-2, using disease-relevant human airway epithelium models and appropriate animal models to assess bioavailability and antiviral efficacy, the mouse model for RSV and the ferret model for SARS-CoV-2. Treatment was considered efficacious when statistically significant reduction in virus titers in nasal lavages (ferrets) and respiratory tissues (mice and ferrets) was observed. Efficacy and cytotoxicity of the drug candidate in multiple cell lines including human airway epithelium were assessed by four-parameter variable-slope regression modeling of 50% inhibitory and cytotoxic concentrations. Timeline endpoints were predefined before initiation of experiments. The number of animal per group (3 to 5, as specified in each figure legend) in *in vivo* efficacy experiments has been selected to provide sufficient statistical power to detect a significant biological effect such as a reduction in viral titers of at least one order of magnitude, based on our previous experience with these models. Before initiation of each individual study, animals were randomly assigned to treatment and control groups. No blinding was used. Primary numerical data are shown in Data S2.

#### Experimental design of mice experiments

6-8–week old female Balb/cJ mice (Jackson laboratory, cat# 000651) were housed in an ABSL-2 facility and rested for 4-5 days. For efficacy studies, mice were randomly divided into groups (n=5) and infected intranasally with $5\times10^5$ $TCID_{50}$ (25 µl per nare) of recRSV-A2line19F-[mKate] in PBS while under anesthesia with ketamine/xylazine. Treatment (4'-FlU or vehicle) was administrated at the indicated time post-infection via oral gavage in a 200 µl suspension of 0.5% Tween80 in 10 mM sodium citrate in water. Temperature and food consumption were monitored daily, body weight was determined twice daily. All animals were euthanized at 4.5 days after infection and lungs were harvested. To determine lung viral titers, lungs were weighted and homogenized with a bead beater in 300 µl PBS in 3 bursts of 20 seconds by 5-minute rest on ice after each cycle. Samples were clarified for 5 minutes at 4°C and 20,000×g, supernatant aliquoted and stored at -80°C before being titrated by median tissue culture infectious dose ($TCID_{50}$) normalized per gram of lung tissue and per ml of lysate. For *in vivo* live imaging, mice were infected with recRSV-A2line19F-[redFirefly] in PBS and treatment was initiated at the indicated time. Bioluminescence was monitored once daily at the indicated time after isoflurane anesthesia and retro-orbital injection with 100 µl of 100 mg/ml D-luciferin (Goldbio). Acquisition was initiated 30s after substrate injection in an IVIS Spectrum (Caliper LifeSciences). Acquisition parameters were a sequence of 9×30-second exposures with medium binning (binning=8), small aperture (f=1) and large field of view (D - 22) using Living Image 4.5.4 Software for Windows 10. For tolerability study, mice were infected with 300,000 $TCID_{50}$ recRSV-A2line19F-[mKate] or mock and treated with 5mg/kg mice body weight 4'-FlU at 12 hours post-infection once daily for 4 days. Blood samples were collected prior to infection and at 1.5, 2.5, 3.5, and 4.5 days post-infection. Complete blood counts were performed with a VETSCAN HM5 (Abaxis) following the manufacturer's protocol.

#### Experimental design of ferret experiments

6-10–month old female ferrets (*Mustela putorius furo*; Triple F Farms) were used as an *in vivo* model to examine the therapeutic efficacy of orally administered 4'-FlU against SARS-CoV-2 infection. Group sizes of 3-4 ferrets were used for efficacy studies. Animals were randomly assigned to the different study groups. No blinding was performed. Viruses were administered to animals through intranasal inoculation. Ferrets were inoculated with SARS-CoV-2 ($1 \times 10^5$ pfu of 2019-nCoV/USA-WA1/2020 or hCoV-19/USA/CA_CDC_5574/2020) in 1 ml (0.5 ml per nare). At 12 hours after infection, a group of ferrets was treated once daily (*q.d.*) with vehicle (10 mM sodium citrate with 0.5% (v/v) Tween 80) or 4'-FlU at a dosage of 20 mg $kg^{-1}$, respectively. Nasal lavages were collected every 12 hours for all ferrets. Once daily dosing was continued for 4 days post infection. All animals were euthanized 4 days after the infection was started. Organs and tissues were harvested and stored at -80°C until processed.
For virus titration, samples were weighed and homogenized in sterile PBS. Tissue homogenates were clarified by centrifugation (2,000×g for 5 minutes at 4°C). The clarified supernatants were then harvested and used in plaque assays. For detection of viral RNA, total RNA was extracted from organs using a RNeasy mini kit (Qiagen), in accordance with the manufacturer's protocol. Total RNA was extracted from nasal lavages using a ZR viral RNA kit (Zymo Research) in accordance with the manufacturer's protocols. Virus titers were determined by plaque assays and viral RNA copy numbers were determined by RT–qPCR quantitation.

**SARS-CoV-2 RNA copy numbers**
Detection of SARS-CoV-2 RNA was performed using the nCoV_IP2 primer–probe set (National Reference Center for Respiratory Viruses, Pasteur Institute). RT–qPCR reactions were performed on an Applied Biosystems 7500 real-time PCR system using the StepOnePlus real-time PCR system. Viral RNA was detected using the nCoV_IP2 primer–probe set in combination with TaqMan fast virus 1-step master mix (Thermo Fisher Scientific). Viral RNA copy numbers were determined based on a standard curve created using a PCR fragment (nucleotides 12669–14146 of the SARS-CoV-2 genome) as previously described. The RNA values were normalized to the weights of the tissues used.

**Institutional Animal Care and Use Committee (IACUC) approval statement.**
All animal work was performed at Georgia State University in compliance with the Guide for the Care and Use of Laboratory Animals of the National Institutes of Health. Mouse work was approved by the GSU Institutional Animal Use and Care Committee (IACUC) under protocols A17019 and A20012, ferret work was approved under protocol A20031.

**Cells**
African green monkey kidney cells VeroE6 (ATCC® CRL-1586™), Vero/hSLAM (expressing human signaling lymphocytic activation molecule), Madin-Darby canine kidney cells (MDCK, ATCC® CCL-34™), human lung adenocarcinoma epithelial cells Calu-3 (ATCC® HTB-55™), human epithelial/HeLa contaminant HEp-2 cells (ATCC® CCL-23™), human bronchial epithelial BEAS-2B (ATCC® CRL-9609™) and baby hamster kidney cells (BHK-21; ATCC® CCL-10™) stably expressing either T7 polymerase (BSR-T7/5) or G protein of rabies virus strain SAD-B19 (BSR-RVG) were cultivated at 37°C and 5% $CO_2$ in Dulbecco's Modified Eagle's medium (DMEM) supplemented with 7.5% fetal bovine serum (FBS). Insect cells from Spodoptera Frugiperda (SF9, ATCC® CRL-1711™) were propagated in suspension using Sf-900 II serum-free media (SFM) (Thermo Scientific) at 28°C. All cell lines used in this study are

routinely checked for mycoplasma and microbial contamination. Mammalian cell transfections were performed using GeneJuice transfection reagent (Invitrogen) while insect cell transfections were performed using Cellfectin II transfection reagent (Gibco). All cell lines were routinely checked for mycoplasma contamination. Normal primary human bronchial/tracheal epithelial cells (HBTEC) from a 30-year old female (LifeLine Cell Technology, cat# LM-0050, lot# 3123, passage 2, "donor F1") were grown in BronchiaLife cell culture medium (LifeLine Cell Technology). Normal human bronchial/tracheal epithelial cells (NHBE) (Lonza Bioscience, cat# CC-2540S, lot# 0000646466, passage 2, donor "M4") from a 38 year-old male were cultured in PneumaCult-Ex Plus (Stemcell Technologies cat# 05040) following the manufacturer's instructions.

**Plasmids**

Plasmids to rescue recombinant RSV A2 with line19 F harboring a mKate reporter (labelled here as recRSV-A2line19F-[mKate]) or RSV A2 with line19 F with hyperfusogenic F_D489E substitution and FireSMASh reporter (labelled here as recRSV-A2line19F-[FireSmash]) were previously described. To rescue a recombinant RSV harboring a red-shifted firefly luciferase (labelled as recRSV-A2line19F-[redFirefly]), we swapped the mKate reporter of the pSynkRSV-line19F vector with the redFirefly sequence in a succession of steps: 1) We fused immediately downstream of the redFirefly sequence from pCMV-Red Firefly (ThermoFisher Scientific cat. 16156) a sequence containing the RSV L noncoding region, L gene end, NS1/NS2 intergenic junction with a BlpI restriction site, mimicking the sequence downstream of mKate on the pSynkRSV-line19F vector. The assembly was used with NEBuilder (New England Biolabs) and the primer sets 5'- GCGGCCGCAAAATCAGCC and 5'-TCATCACATCTTGGCCACGG for pCMV-Red Firefly and with 5'-ccgtggccaagatgtgatgaGTATTCAATTATAGTTATTAAAAACTTAACAG) and 5'-gaggctgattttgcggccgcGCTAAGCAAGGGAGTTAAATTTAAG for the downstream sequence with BlpI. 2) The sequence of redFirely and its downstream sequence up to the BlpI site were amplified by PCR to insert a second BlpI site immediately upstream of the redFirefly sequence using primers 5'-catcatGCTTAGCatggaaaatatggaaaacg and 5'-gaggctgattttgcggccgcGCTAAGCAAGGGAGTTAAATTTAAG, mimicking the sequence upstream of mKate on the pSynkRSV-line19F vector. The amplified product was digested with BlpI (New England Biolabs) and purified. 3) Since BlpI is not a unique site on the pSynkRSV-line19F vector, we generated an intermediate shuttle vector by inserting a fragment from pSynkRSV-line19F vector encompassing the leader promoter sequence to the nucleoprotein gene and containing mKate, by digesting it with AvrII (New England Biolabs) and PmeI (New England Biolabs). In parallel we digested and purified the backbone of a pCDNA 3.1 (ThermoFisher Scientific cat. V79020) with AvrII (New England Biolabs) and PmeI (New England Biolabs), and the two fragments were ligated with the t4 ligase ((New England Biolabs). 4) The shuttle vector was opened with BlpI to release the mKate segment and the backbone purified, and ligated to the purified amplicon of step (2) containing redFirefly. 5) The shuttle vector with redFirefly was opened with AvrI and PmeI and the insert ligated back to the backbone of the pSynkRSV-line19F vector. The final construct was confirmed with Sanger sequencing. For expression of SARS-CoV-2 nsp12, nsp8 and nsp7, SARS-CoV-2 RNA was isolated from stocks of isolate USA-WA1/2020 (BEI# NR-52281) and reverse transcribed using superscript III (Invitrogen). The resulting cDNA was used as a template for the amplification of nsp7, nsp8 and nsp12 encoding genes the primer 5'-

CTGGACATATGGGCAGCAGCCATCATCATCATCATCACAGCAGCGGCGAAAACCTG TATTTTCAGGGCTCTAAAATGTCAGATGTAAAG and 5’-CTAGAGCGGCCGCCTATTGTAAGGTTGCCCTGTTG (nsp7), 5’-CTGGATCTAGAAATAATTTTGTTTAACTTTAAGAAGGAGATATAATGGGCAGCAGCC ATCATCATCATCAT and 5’-CTAGAGCGGCCGCCTACTGTAATTTGACAGCAGAATTG (nsp8), and 5’-CTGGACATATGTCAGCTGATGCACAATCG and 5’-CTAGACTCGAGGCCCTGAAAATACAGGTTTTCGCCGCTGCTCTGTAAGACTGTATGC GGTG (nsp12). PCR amplified products were individually cloned into pET30a(+) expression vector (Novagen) with tobacco etch virus protease-cleavable 6×His tag at the N-terminus of nsp7 and nsp8, and at the C-terminus of nsp12 using appropriate restriction sites and cloning strategies, bringing the cloned genes under T7 promoter control. A catalytically inactive variant of nsp12 was generated by site-directed mutagenesis, modifying residues 762-SDD in the conserved catalytic motif C to 762-SNN with appropriately designed primers.

**Viruses**

Recombinant RSVs were rescued as described previously, briefly BSR-T7/5 cells were co-transfected with the cDNA genome along with helper plasmids encoding the RSV L, N, P and M2-1 protein, further amplified on HEp-2 cells after a freeze-thaw cycle and supernatant clarification (1800×g, 5 minutes, 4°C). Stocks were prepared as described previously by infecting 50% confluent HEp-2 cells in 15cm dishes at a multiplicity of infection (MOI) of 0.01 $TCID_{50}$/cell, incubated at 37°C overnight then transferred to 32°C for five days. Infected cells were scraped, virions released with a freeze/thaw cycle and supernatant clarified (1800×g, 5 minutes, 4°C). Screening-grade stocks of recRSV-A2line19F-[FireSmash]) were grown in the presence of 3 μM asunaprevir to reduce accumulation of Firefly luciferase and further purified at the interface of a 20%/60% sucrose gradient in TNE buffer (50 mM Tris/Cl pH 7.2, 10 mM EDTA) with centrifugation at 30,000rpm on a SW41 rotor (Beckman Coulter) for 2 hours at 4°C. Clinical RSV B isolates 16F10 (GenBank: KY674983.1) and 6A8 (GenBank: MF001044.1) were a kind gift from A. Greninger. Samples were collected from patient nasal wash specimens in 2017, cultured on primary rhesus monkey kidney (RhMK) cells, and amplified once on HEp-2 cells prior to use in this study. RSV viral titers were determined using standard 50% tissue infective dose ($TCID_{50}$) assay in HEp-2 cells and 96 well plates, with a Spearman and Karber based method using either fluorescence or immunostaining for detection. Recombinant measles virus strain Edmonston with nano luciferase reporter including a destabilizing PEST sequence (labelled here as recMeV-[$N_{PEST}$luc]) was rescued and amplified on Vero/hSLAM cells. Recombinant respiroviruses human parainfluenza 3 virus with Nano luciferase reporter (HPIV3-JS NanoLuc, labelled here recHPIV-3-[Nluc]), Sendai virus with Gaussia luciferase reporter labelled here recSeV-[Gluc] were rescued and amplified on VeroE6 cells. Recombinant rhabdoviruses, vesicular stomatitis virus (Indiana strain) with Nano luciferase reporter (labelled here as recVSV-[Nluc]) and rabies virus strain SAD-B19 with G deletion and Nano luciferase reporter (labelled here as recRabVv-[Nluc]) were rescued and authenticated through RT-PCR and Sanger-sequencing. SARS-CoV-2 isolates USA-WA1/2020 (BEI# NR-52281) and USA/CA_CDC_5574/2020 (BEI# NR-54011) were sequence-validated and propagated on VeroE6 and Calu-3 cells, in DMEM supplemented with 2% FBS following approved procedures in biosafety-level-3 containment.

**Minireplicon assays**

For RSV-derived minireplicon assays, a plasmid expressing the RSV minigenome containing the firefly luciferase reporter, under control of RNA pol I promoter, and a set of helper plasmids expressing codon-optimized RSV P, L, N and M2-1 proteins, under the control of CMV promoter, were co-transfected with GeneJuice reagent (Millipore Sigma) following manufacturer's instructions in 50% confluent HEK-293T cells. For dose-response experiments, cells were transfected in 96-well plate format. 3-fold serial dilutions of compounds 4'-FlU or NHC were prepared in triplicate using a Nimbus liquid handler (Hamilton) and transferred on transfected cells 4 hours post-transfection. At 48 hours post-transfection, luciferase activities were determined using ONE-Glo luciferase substrate (Promega) and a H1 synergy plate reader (BioTek). Each plate contained 4 wells of each positive and negative controls (minireplicon with media containing dimethyl sulfoxide (DMSO) or minireplicon with L-expressing plasmid replaced by pCDNA3.1). Normalized luciferase activities were analyzed with the formula: % inhibition = $(Signal_{Sample}-Signal_{Min})/(Signal_{Max}-Signal_{Min})\times100$, and dose response curves were further analyzed by normalized non-linear regression with variable slope to determine 50% effective concentration ($EC_{50}$) and 95% CIs Prism 9.0.1 for MacOS (GraphPad). Similarly, luciferase activities from MeV strain Edmonston-based minireplicon (firefly luciferase), NiV minireplicon (Nano luciferase) and the recHPIV3-JS-NanoLuc (Nano luciferase, labelled in this context as HPIV3 maxireplicon), were transfected in BSR-T7/5 as previously described.

**Dose response antiviral assays**

For reporter-based dose-response assays, 3-fold serial dilutions of 4'-FlU or NHC were prepared in triplicate using a Nimbus liquid handler (Hamilton) and transferred to 96-well plates seeded the day before at 50% confluence in 96-well plate format. Immediately after addition of compound, cells were infected with either recRSV-A2line19F-[FireSmash], recMeV-[NPESTluc] recHPIV-3-[Nluc], recSeV-[Gluc], recVSV-[Nluc], or recRabV-[Nluc] at MOI 0.2 $TCID_{50}$/cell. At 48 hours post-transfection, luciferase activities of reporter-expressing viruses were determined using either ONE-Glo luciferase substrate, Nano-Glo Dual-Luciferase substrate (Promega) and a H1 synergy plate reader (Biotek). Each plate contained 4 wells each of positive and negative control (infected cells with media containing DMSO or 100 µM cycloheximide, respectively). Normalized luciferase activities were analyzed with the formula: % inhibition = $(Signal_{Sample}-Signal_{Min})/(Signal_{Max}-Signal_{Min})\times100$, and dose response curves were further analyzed by normalized non-linear regression with variable slope to determine 50% effective concentration ($EC_{50}$) and 95% CIs with Prism 9.0.1 for MacOS (GraphPad). For RSV virus yield reduction, HEp-2 cells were infected with recRSV-A2line19F-[mKate] or clinical isolates in 12-well plate format at MOI 0.1 $TCID_{50}$/cell for 2 hours at 37°C. Inoculum was removed and replaced with DMEM with 2% FBS and indicated concentrations of compound and cells were incubated for 3 days. Viral titers were determined by standard $TCID_{50}$ with fluorescence or immunostaining for detection. For SARS-CoV-2 virus yield reduction on VeroE6 cells, cells were seeded in 12-well plates (300,000 cells per well) the day before infection, infected at a MOI of 0.1 PFU/cell with a 1 hour absorption step and inoculum was removed and replaced with fresh DMEM with 2% FBS and indicated concentrations of 4'-FlU (vehicle: 0.1% DMSO). Infected cells were incubated with compound for 48 hours at 37 °C, followed by virus titration by standard plaque assay. Log viral titers were normalized using the average top plateau of viral titers to define 100% and were analyzed with a non-linear regression with variable slope to determine $EC_{50}$ and 95% CIs with Prism 9.0.1 for MacOS (Graphpad).

**Cytotoxicity assays**

To determine the effect of compound on cell metabolism, HEp-2, MDCK, Beas-2B, or BHK-T7 cells were seeded at 50% confluence in 96-well plates and were incubated with 3-fold serial dilution of 4'-FlU from 500 µM as described for dose-response assays, including positive and negative controls for normalization. After 48-hour incubation at 37°C, cells were incubated with PrestoBlue (ThermoFisher Scientific) for 1 hour at 37°C and fluorescence measured with a H1 synergy plate reader (Biotek). 50% cytotoxic concentrations ($CC_{50}$) and 95% CIs after normalized non-linear regression and variable slope were determined using Prism 9.0.1 for MacOS (GraphPad). To uncover potential mitochondrial toxicity masked by the Crabtree effect, BEAS-2B cells were also incubated in the presence of glucose-free RPMI media supplemented with galactose as carbohydrate source. To further assess the potential inhibitory effect on mitochondrial and nuclear polymerases, HBTEC cells were seeded at 50% confluence in 96 well plates and incubated with 3-fold dilutions of compound, and intracellular concentration of two mitochondrial proteins, the mitochondrial DNA-encoded COX-I and the nuclear DNA-encoded SDH-A was determined with the in-cell ELISA Mitobiogenesis kit following the manufacturer's instructions (Abcam cat# ab110217).

**Ribonucleotide competition of RSV inhibition**

HEp-2 cells were infected with RSV-A2line19F-[FireSMASh] at a MOI of 0.1 $TCID_{50}$/cell, maintenance media was supplemented with 4'-FlU at 10 µM alone or in combination with 0.1 to 300 µM exogenous ribonucleosides (Sigma-Aldrich). Firefly luciferase reporter activity was quantified at 48 hours post-infection. Values are expressed relative to the values for the vehicle-treated samples.

**Recombinant RSV L and P protein production and purification**

For purification of RSV L and P proteins (A2 strain), a pFastBac Dual plasmid (Invitrogen) containing the codon-optimized open reading frames of L and P proteins under the control of polyhedrin and p10 promoters, respectively, was used to recover recombinant baculoviruses using the Bac-to-Bac Baculovirus Expression System and SF9 cells. The P protein sequence contains a C-terminal 6-histidines tag separated by a tobacco etch virus (TEV) cleavage site, allowing co-purification of L-P complexes by immobilized metal affinity chromatography (IMAC). SF9 cells were infected in suspension with MOI 1 PFU/cell for 72-80 hours, pelleted and gently lysed on ice for 45 minutes with a buffer containing 50 mM $NaH_2PO_4$ [pH 8.0], 150 mM NaCl, 20 mM imidazole, 0.5% NP-40 with Pierce Protease Inhibitor and Pierce universal nuclease (ThermoFisher Scientific). Clarified lysates (30 minutes, 15,000×g, 4°C) were incubated for 2 hours with shaking with pre-equilibrated HisPur Ni-NTA Resin (ThermoFisher Scientific). After 5 washes with 10 bed volumes of lysis buffer with 60 mM imidazole, proteins were eluted with 3 bed volumes of lysis buffer with 250 mM imidazole. Eluates were dialyzed overnight in storage buffer: 20 mM Tris-HCl [pH 7.4], 150 mM NaCl, 10% glycerol, 1 mM dithiothreitol, aliquoted and stored at -80°C. Uncropped SDS-PAGE scans are available in Data S1.

**Recombinant SARS-CoV-2 nsp7, nsp8, and nsp12 protein production and purification**

The sequence confirmed constructs of nsp7 and nsp8 were transformed into BL21(DE3)pLysS cells. For the expression of nsp12 in *E. coli*, first chaperon plasmid pG-Tf2 (Takara Biosciences) was transformed into BL21 cells to enhance correct folding and solubility of recombinant proteins, followed by transformation with pET30a-nsp12 plasmid. The transformed cells were grown in Luria-Bertani (LB) broth for nsp7 and nsp8 at 37°C until the $OD_{600}$ reached 0.6. nsp7 and nsp8 cultures were induced with 0.5 mM isopropyl β-d-1-thiogalactopyranoside (IPTG) (Teknova) and the temperature was reduced to 18°C. The expression culture was grown for ~16 hours. nsp12 expression was performed by growing the cells in Terrific broth (TB) at 37°C until the $OD_{600}$ reached 0.6. Tetracycline was added to a final concentration of 10 ng/ml and the temperature was reduced to 18°C. After half an hour incubation, 0.5 mM IPTG was added to the culture and incubated at 18°C for ~16 hours. Cells were harvested by centrifugation and the cell pellet was stored at -20°C until further use. The cell pellet was resuspended in lysis buffer (Tris 50 mM pH 8.0, NaCl 200 mM, glycerol 10%, 20 mM imidazole, protease inhibitor cocktail (Pierce), Pierce universal nuclease (ThermoFisher Scientific)) with 0.3mg/ml lysozyme and incubated on ice for ~20 minutes. The samples were sonicated for ~6-8 minutes. Cell debris were separated by centrifugation at 14,000×g for 30 minutes, clarified supernatant collected, and loaded on to HisPur Ni-NTA resin (ThermoFisher Scientific) pre-equilibrated with lysis buffer and incubated for ~45 minutes at 4°C. Beads were washed extensively with wash buffer (Tris 50 mM pH 8.0, NaCl 200 mM, 10% glycerol containing 20-100 mM imidazole), and proteins eluted with Tris 50 mM pH 8.0, NaCl 200 mM, glycerol 10% containing 250 mM imidazole. The purified fractions were pooled, and the histidine tag was cleaved by adding TEV protease to the purified protein which was also simultaneously dialyzed overnight against the 1 L dialysis buffer (Tris 30 mM pH 7.4, NaCl 150 mM, glycerol 10%). Dialyzed 6×His tag cleaved proteins were re-incubated with HisPur Ni-NTA resin pre-equilibrated with Tris 50 mM pH 7.4, NaCl 200 mM, glycerol 10%, 20mM imidazole for 15 minutes. Flow through containing the tag cleaved proteins was collected and the resin was further washed with buffer containing imidazole. The desired His tag cleaved fractions were pooled and dialyzed against storage buffer (Tris 25 mM pH 7.4, NaCl 150 mM, glycerol 10%, DTT 1 mM) at 4°C overnight. The proteins were concentrated to final concentrations of 60, 30 and 6 µM for nsp7, nsp8 and nsp12 respectively, flash frozen and stored at -80°C for further use. Two additional polypeptides that co-purified with nsp12 likely represent host Gro-ES (60 kDa) and trigger factor (Tf) (56 kDa) proteins. Uncropped SDS-PAGE scans are available in Data S1.

#### *In vitro* RNA synthesis assays

3’ primer-extension assays were performed based on an established assay with modifications. Briefly, purified polymerase complexes (100-200 ng of L protein) were incubated in 5 µl final volume with a transcription buffer containing 20 mM Tris-HCl pH 7.4 [RT], 30 mM NaCl, 10% glycerol, 1 mM dithiothreitol, 8 mM $MgCl_2$, 4 µM RNA template, 20 µM RNA primer, indicated concentrations of nucleotides and 1 µCi of (alpha)32P-labelled ATP (Perkin-Elmer). HPLC-purified 5’ phosphorylated RNA templates and non-phosphorylated RNA primers (SIGMA-ALDRICH) were used to reduce non-specific extension of the template. Reactions were performed for 1 hour at 30°C and the reaction was stopped with 1 volume of deionized formamide with 25 mM ethylenediaminetetraacetic acid (EDTA). Following 5 minutes denaturation at 95 °C, samples were separated by 7M urea Tris-Borate-EDTA 20% polyacrylamide gel electrophoresis and visualized by autoradiography with X-ray films (CL-XPosure, ThermoFisher Scientific) or with a storage phosphor screen BAS IP MS 2040 E (GE Healthcare Life Sciences) and imaged with Typhoon FLA 7000 (GE Healthcare Life Sciences). Densitometry analysis was performed using FIJI 2.1.0. For quantifications of nucleotide incorporation kinetics, the intensity of each band corresponding to a primer-extension product was individually measured for each lane, and either the band corresponding to the UTP incorporation or the pooled primer-extension products following initial UTP incorporation were normalized to the pooled intensities of all the primer-extension products in that lane. Enzyme kinetics ($V_{max}$ and Km) were determined using the Michaelis-Menten equation with Prism 9.0.0 for macOS (GraphPad). Nucleoside selectivity was determined using the nucleotide analog/nucleotide ratio of each Vmax/Km ratios. For side-by-side comparison of elongation products, reactions were performed as previously noted with saturating concentration of 300 µM of UTP or 4'-FlU-TP. Each band corresponding to a primer-extended product containing 4'-FlU-TP was directly normalized to the equivalent UTP-containing band. For *de novo* RNA synthesis, an RNA template corresponding to the 25 nt of the RSV trailer complement sequence was incubated at 2 µM with RSV RdRP (100-200 ng of L), 8 mM $MgCl_2$, 1 mM dithiothreitol, indicated concentrations of each nucleotides and 10 µCi of (alpha)32P-labelled GTP (Perkin-Elmer), 20 mM Tris-HCl [pH 7.4], 15 mM NaCl, 10 % glycerol. Reactions were equilibrated 5 minutes at 30°C before addition of RSV RdRP, then incubated at 30°C for 3 hours. RNAs were precipitated overnight at -20°C with 2.5 volumes of ethanol, 0.1 volume of 3M sodium acetate plus 625 ng of glycogen (ThermoFisher Scientific). Pellets were washed with 75% ethanol, dried and resuspended in 10 µl of 50% deionized formamide. After 5 minutes denaturation at 95°C, RNAs were separated as described previously and visualized by autoradiography using CL-XPosure™ Film (ThermoFisher Scientific). SARS-CoV-2 polymerase *in vitro* assays were set up analogous to the protocol described above. All three separately purified proteins nsp7 (6 µM), nsp8 (3 µM) and nsp12 (300 nM) were incubated in 20 mM Tris-HCl pH 7.4, 10% glycerol, 1 mM dithiothreitol, 5 mM $MgCl_2$, 1 µM RNA template, 20 µM RNA primer, 20 nM ATP, 20 nM CTP and 1 µCi of (alpha)32P-labelled ATP (Perkin-Elmer). Primer-template sets were used as described above. For the comparison of primer extension reaction 200 nM of UTP or 4'-FlU-TP was used. Uncropped autoradiograms including biological repeats are available in Data S1.

**Cellular uptake and anabolism**

$10^5$ HBTECs were seeded per well in 24-well plates. The next day the media was supplemented with 20 µM 4'-FlU or vehicle (DMSO). For wash-in studies, cell culture media was removed at the indicated time-point and cells were washed twice with Dulbecco's modified phosphate buffered saline without calcium and magnesium (DPBS) and harvested using 500 µl of ice-cold 70% methanol. Cellular extracts were clarified by centrifugation at 16,000×g for 10 minutes at 4°C and stored at -20°C until further analysis. For wash-out studies, cells were incubated in the presence of 20 µM 4'-FlU or vehicle (DMSO) for 24 hours, then washed twice with DPBS and incubated with maintenance media for the indicated time point before harvesting. 4'-FlU and 4'-FlU-TP anabolite were quantitated by qualified internal standard-based LC-MS/MS method using an Agilent 1200 system (Agilent Technologies) equipped with a SeQuant ZIC-pHILIC column (The Nest Group). MS analysis was performed on a QTrap 5500 mass spectrometer (AB Sciex) using negative-mode electrospray ionization (ESI) in the multiple-reaction-monitoring (MRM) mode. Data analysis was done with Analyst software (AB Sciex).

**Air-liquid interfaces of primary human airway epithelial (HAE) cells**

30,000 HBTEC or NHBE cells at passage 2 were seeded on 6 mm 0.4 µm pore size polyester inserts (Corning Costar Transwell) and differentiated at an air-liquid interface using Air-Liquid Interface Differentiation Medium (Lifeline cell technology, cat# LL-0023) for the former or Pneumacult-ALI (Stemcell Technologies cat# 05001) according to the manufacturer's instruction. Transepithelial electrical resistance was monitored using the EVOM instrument and STX2 electrode (World Precision Instrument).

**Apical shed viral titer determination**

Differentiated primary cells were washed on the apical side with phosphate-buffered saline (PBS) without calcium and magnesium. Cells were infected apically either with recRSV-A2line19F-[mKate] (500,000 $TCID_{50}$) or SARS-CoV-2 for (40,000 PFU) for 2 hours at 37°C. Compound was added in the basal media at the indicated concentrations and 0.1% final DMSO. To harvest shed virus, ALI-HAE cells were incubated apically with 200 µl Dulbecco's phosphate-buffered saline without calcium and magnesium at 37°C for 30 minutes, aliquoted and stored at -80°C before titration. For RSV, shed viruses were harvested at peak at day 3 post-infection, and tittered by standard $TCID_{50}$. For SARS-CoV-2, samples were harvested at 24, 48, 72 and 96 hours post-infection ("donor F1") or 24, 48, 72 and 144 hours post-infection ("donor M4"), For SARS-CoV-2 titration, VeroE6 cells were seeded in 12-well plates at 200,000 cells per well. The next day, samples were thawed and serially diluted three times with 10-fold dilutions in DMEM. VeroE6 cell-culture media was removed and cells were incubated with 100 µl of diluted inoculum with rocking every 10 minutes for 1 hour at 37°C. The overlay medium (DMEM+2% FBS+1.2% microcrystalline cellulose Avicel 581-NF (FMC)) was added on the cells and the plates were incubated for three days at 37°C. The plaques were revealed after two washes with PBS, 1-hour fixation with 10% neutral buffered formalin, and 15 minutes coloration with a 1% crystal violet solution in 20% ethanol.

**Antibodies**

For immunostaining of infected cells in virus titer determination, fixed and permeabilized cells were successively stained with goat anti-RSV 1:1000 (Millipore Sigma, cat. AB1128) followed by donkey anti-goat HRP-coupled 1:1000 (Jackson Immunoresearch cat. 705-035-147) and infected cells were visualized with Trueblue peroxidase substrate according to the manufacturer's instructions (KPL). For RSV confocal microscopy experiments, fixed and permeabilized cells were stained with mouse anti-(human) ZO-1 1:50 (BD Biosciences, cat. 610966), goat anti-RSV 1:1000 (Millipore Sigma, cat. AB1128), rabbit anti-beta IV Tubulin recombinant antibody Alexa Fluor® 647-conjugated [EPR16775] 1:100 (Abcam, cat. ab204034) followed by donkey anti-goat Alexa Fluor 568 1:500 (ThermoFisher Scientific, cat. A-11057) and rabbit anti-mouse IgG (H+L) cross-adsorbed secondary antibody, Alexa Fluor 488 1:500 (ThermoFisher Scientific, cat. A1059). For SARS-CoV-2 confocal microscopy, cells were stained with rabbit anti-SARS-CoV-2 Nucleocapsid monoclonal antibody (HL453) (Invitrogen, MA5-36272) 1:100 and mouse anti-MUC5AC 1:200 (ThermoFisher MA5-12175), followed by donkey anti-goat Alexa Fluor 568 1:500 (ThermoFisher Scientific, cat. A-11057) and rabbit anti-mouse IgG (H+L) Cross-Adsorbed Secondary Antibody, Alexa Fluor 488 1:500 (ThermoFisher Scientific, cat. A-11059).

**Confocal microscopy**

Differentiated human airway epithelial cells were either infected (or mock-infected) with recRSV-A2line19F-[FireSmash] (500,000 $TCID_{50}$), washed with PBS daily and fixed at 5 days post-infection with 200 µl of 4% paraformaldehyde in PBS for 1 hour at RT, or were infected (or mock-infected) with 40,000 plaque-forming units of SARS-CoV-2 isolate USA-WA1/2020 (BEI# NR-52281) and fixed at day 3 post-infection. Cells were permeabilized for 20 minutes with PBS with 0.1% Triton X-100 and 3% bovine serum albumin (BSA). After three 5-minute washes with wash buffer (PBS with 0.3% BSA), cells were incubated with 100 µl of primary antibody in wash buffer for 1 hour at RT. After 3×5 minutes 200 µl washes with wash buffer, cells were incubated with secondary antibody in wash buffer for 1hour at RT. Cells were washed and incubated with Hoechst 34580 at 1/1000 in wash buffer for 5 minutes. After three 5-minute washes with wash buffer, membranes were cut and mounted between glass slides and coverslips using Prolong Diamond antifade mountant (ThermoFisher Scientific) and edges were sealed with nail polish. Image capture were performed with a Zeiss LSM 800 confocal microscope coupled with an Airyscan module and the Zeiss Zen Blue software.

**Statistical analysis**
Source data for all numerical assays conducted in this study are provided in Data S2. Excel and GraphPad Prism software packages were used for data analysis. One-way and two-way ANOVA with Dunnett's or Tukey's multiple comparisons post-hoc tests without further adjustments - as specified in figure legends - were used to evaluate statistical significance when more than two groups or two parameters were compared. The specific statistical test applied to individual studies is specified in figure legends. When calculating antiviral potency and cytotoxicity, effective concentrations were calculated from dose-response data sets through 4-parameter variable slope regression modeling, and values are expressed with 95% CIs when available. Biological repeat refers to measurements taken from distinct samples, and results obtained for each biological repeat are shown in the figures along with the exact sample size (n). For all experiments, the statistical significance level alpha was set to <0.05, exact P values are shown in individual graphs.

**Synthetic chemistry**
5'-Deoxy-5'-iodouridine (Fig. S16). A 1 L three-necked round-bottomed flask flushed with argon and fitted with a thermometer and addition funnel was charged with uridine (35 g, 143 mmol), triphenylphosphine (56.4 g, 215 mmol), imidazole (14.64 g, 215 mmol), and anhydrous THF (400 ml). The suspension was stirred vigorously for 30 minutes while cooling to 0°C and then treated with a THF (100 ml) solution of iodine (40 g, 157.7 mmol) dropwise over a 2-hour period. The mixture was warmed to room temperature. After 16 hours, tlc (10% methanol in methylene chloride) indicated complete consumption of starting uridine. The mixture was concentrated under vacuum followed by a solvent exchange with isopropanol (400 ml). Upon cooling with an ice-bath the product precipitated out of solution as a white solid which was collected by vacuum filtration and washed with ice-cold isopropanol (150 ml) followed by hexanes (100 ml) to give 5'-deoxy-5'-uridine (35 g, 69% yield).
$^1$H NMR (400 MHz, DMSO-$d_6$) δ 11.40 (s, 1H), 7.68 (d, $J$ = 8.1 Hz, 1H), 5.80 (d, $J$ = 5.9 Hz, 1H), 5.68 (dd, $J$ = 8.0, 2.2 Hz, 1H), 5.50 (d, $J$ = 5.8 Hz, 1H), 5.38 (d, $J$ = 5.0 Hz, 1H), 4.19 (q, $J$ = 5.7 Hz, 1H), 3.86 (dq, $J$ = 14.0, 4.5 Hz, 2H), 3.55 (dd, $J$ = 10.5, 5.4 Hz, 1H), 3.40 (dd, $J$ = 10.5, 6.5 Hz, 1H).
4',5'-Didehydro-5'-deoxyuridine (Fig. S16). In a 1 L round-bottomed flask, a suspension of 5'-deoxy-5'-iodouridine (34.55 g, 226 mmol) in methanol (350 ml) was treated with sodium methoxide (67.3 ml, 293 mmol). Upon heating to 60°C the mixture became homogeneous. After 3.5 hours, the mixture was cooled to room temperature and acidified to pH 7 by treating with dry ice. The mixture was filtered, concentrated under vacuum and then purified by column

chromatography over silica gel using a mobile phase of 10% methanol in methylene chloride to give 4',5'-Didehydro-5'-deoxyuridine (11.2 g, 51% yield) as a white solid.

$^1$H NMR (400 MHz, DMSO-$d_6$) δ 11.44 (s, 1H), 7.59 (d, $J$ = 8.1 Hz, 1H), 5.96 (d, $J$ = 5.4 Hz, 1H), 5.64 (d, $J$ = 8.1 Hz, 1H), 5.60 (d, $J$ = 5.8 Hz, 1H), 5.46 (d, $J$ = 5.7 Hz, 1H), 4.38 (t, $J$ = 5.5 Hz, 1H), 4.33 (s, 1H), 4.24 (q, $J$ = 5.5 Hz, 1H), 4.17 (d, $J$ = 1.8 Hz, 1H).

5'-Deoxy-4'-fluoro-5'-iodouridine (Fig. S16). An oven-dried 500 ml round-bottomed flask was charged with 4',5'-didehydro-5'-deoxyuridine (6.55 g, 29 mmol) and anhydrous acetonitrile (60 ml). The suspension was stirred vigorously for 30 minutes, cooled to 0°C under argon and then treated with triethylamine-trihydrofluoride (2.36 ml, 14.5 mmol) followed by the addition of *N*-iodosuccinimide (8.47 g, 37.7 mmol). After 1 hour at 0°C, the mixture was warmed to room temperature for 16 hours and then filtered. The collected solid was washed with methylene chloride (75 ml) followed by ether (75 ml) and then dried under high vacuum to give 5'-deoxy-4'-fluoro-5'-iodouridine (4.2 g, 40%) as an off-white solid.

$^1$H NMR (400 MHz, Methanol-$d_4$) δ 7.77 (d, $J$ = 8.1 Hz, 1H), 6.05 (s, 1H), 5.69 (d, $J$ = 8.1 Hz, 1H), 4.43 (dd, $J$ = 18.2, 6.5 Hz, 1H), 4.25 (d, $J$ = 6.6 Hz, 1H), 3.85 – 3.63 (m, 2H).

$^{19}$F NMR (376 MHz, Methanol-$d_4$) δ -112.49 (ddd, $J$ = 20.9, 18.1, 6.1 Hz).

2',3'-Di-*O*-acetyl-5'-deoxy-4'-fluoro-5'-iodouridine (Fig. S16). A 500 ml round-bottomed flask was charged with 5'-deoxy-5'-iodo-4'-fluorouridine (15 g, 40.3 mmol), 4-dimethylaminopyridine (250 mg, 2.02 mmol),triethylamine (16.8 ml, 120.9 mmol) and methylene chloride (200 ml). The mixture was treated dropwise with acetic anhydride (11.4 ml, 120.9 mmol) while maintaining a reaction temperature below 30°C. After 3 hours at room temperature, the reaction mixture was quenched with saturated sodium bicarbonate solution. The organic layer was separated, washed with water followed by 1N HCl, dried and concentrated under vacuum to give 2',3'-di-*O*-acetyl-5'-deoxy-4'-fluoro-5'-iodouridine (16 g, 87%) as an off-white solid.

$^1$H NMR (400 MHz, DMSO-$d_6$) δ 11.60 (d, $J$ = 2.1 Hz, 1H), 7.77 (d, $J$ = 8.0 Hz, 1H), 6.03 (d, $J$ = 2.4 Hz, 1H), 5.83 – 5.67 (m, 2H), 5.57 (dd, $J$ = 7.4, 2.4 Hz, 1H), 3.63 (dd, $J$ = 11.8, 8.2 Hz, 1H), 3.51 (dd, $J$ = 22.9, 11.8 Hz, 1H), 2.09 (s, 6H).

$^{19}$F NMR (376 MHz, DMSO-$d_6$) δ -105.54 (m).

5'-(3-Chlorobenzoyloxy)-2',3'-di-*O*-acetyl-4'-fluorouridine (Fig. S16). A 100 ml round-bottomed flask was charged with tetrabutylammonium hydrogen sulfate (3.64 g, 10.7 mmol)1-[(4R,6R)-4'-FlUoro-4-(iodomethyl)-2,2-dimethyl-6,6a-dihydro-3aH-furo[3,4-d][1,3]dioxol-6-yl]pyrimidine-2,4-dione (11.7g, 28.39mmol), 3-chlorobenzoic acid (1.76 g, 11.2 mmol), potassium phosphate dibasic (1.68 g, 9.65 mmol) and water (11 ml). After stirring for 20 minutes, the mixture was treated with a methylene chloride (30 ml) solution of 2',3'-di-*O*-acetyl-5'-deoxy-4'-fluoro-5'-iodouridine (2.00 g, 4.38 mmol) followed by the addition of 3-chloroperoxybenzoic acid (4.93 g, 22.0 mmol) in 4 portions over a 30-minute period. The mixture continued to stir at RT for 16 hours. The mixture was periodically treated with additional potassium phosphate dibasic to maintain pH 3.5. The reaction mixture was quenched by addition of sodium sulfite (5 g) in small portions while maintaining a reaction temperature below 30°C. After stirring for an additional 15 minutes, the mixture was filtered through a pad of Celite. The aqueous layer of the filtrate was extracted with methylene chloride (2 × 15 ml), and then combined organic layers were concentrated under vacuum. The resulting residue was redissolved in ethyl acetate (100 ml) and washed with 1N HCl (2 × 50 ml) followed by saturated sodium carbonate solution (2 × 50 ml). The organic layer was concentrated and the resulting solid purified by column chromatography over silica gel (21 mm x 160 mm) using an ethyl acetate/hexane gradient to give 5'-(3-chlorobenzoyloxy)-2',3'-di-*O*-acetyl-4'-fluorouridine (1.1 g, 52% yield) as a white solid.

$^1$H NMR (400 MHz, DMSO-$d_6$) δ 11.59 (s, 1H), 7.94 (d, *J* = 7.7 Hz, 1H), 7.89 (s, 1H), 7.78 (d, *J* = 8.1 Hz, 2H), 7.60 (t, *J* = 7.9 Hz, 1H), 6.10 (s, 1H), 5.94 (dd, *J* = 20.2, 7.4 Hz, 1H), 5.64 (dd, *J* = 30.4, 7.6 Hz, 2H), 4.73 – 4.48 (m, 2H), 2.09 (s, 3H), 2.01 (s, 3H).

$^{19}$F NMR (376 MHz, DMSO-$d_6$) δ -115.94 (dt, *J* = 20.7, 10.6 Hz).

4'-Fluorouridine (6) (Fig. S16). A thick wall round-bottomed pressure vessel was charged with 5'-(3-chlorobenzoyloxy)-2',3'-di-*O*-acetyl-4'-fluorouridine (1.55 g, 3.2 mmol) and 4N ammonia in methanol (4.15 ml, 191.8 mmol). The mixture was stirred at RT for 1.5 hours after which time tlc (10% methanol in methylene chloride) indicated complete consumption of starting material. The mixture was concentrated under vacuum at 19°C, and the resulting residue triturated with methyl *ter*t-butyl ether. The solid was co-evaporated with methanol and then recrystallized from ethanol (15 ml) to give 4'-fluorouridine (300 mg, 36% yield) as a white solid.

$^1$H NMR (400 MHz, DMSO-$d_6$) δ 11.41 (s, 1H), 7.67 (d, *J* = 8.1 Hz, 1H), 5.97 (d, *J* = 2.8 Hz, 1H), 5.66 (d, *J* = 5.4 Hz, 1H), 5.64 (d, *J* = 8.1 Hz, 1H), 5.47 (t, *J* = 5.9 Hz, 1H), 5.16 (d, *J* = 8.8 Hz, 1H), 4.24 (ddd, *J* = 17.7, 8.8, 6.5 Hz, 1H), 4.12 (td, *J* = 6.0, 2.8 Hz, 1H), 3.54 (t, *J* = 5.6 Hz, 2H).

$^{19}$F NMR (376 MHz, DMSO-$d_6$) δ -120.94 (dt, *J* = 17.7, 5.4 Hz).

LCMS Calculated for $C_9 H_{12}FN_2O_6$ [M+H$^+$]: 263.0; found: 263.0

2',3'-Di-*O*-benzyloxycarbonyl-5'-deoxy-4'-fluoro-5'-iodouridine (Fig. S17). A 150 ml round-bottomed flask was charged with 5'-deoxy-5'-iodo-4'-fluorouridine (2.6 g, 6.99 mmol) and methylene chloride (35 ml). After stirring for 20 minutes at RT, the suspension was cooled to 0°C and treated with benzyl chloroformate (4.49 ml, 31.44 mmol) followed by dropwise addition of 1-methylimidazole (3.34 ml, 41.93 mmol) over a 10-minute period. The mixture was stirred an additional 10 minutes at 0°C and then allowed to slowly warm to room temperature. After 18 hours, the turbid mixture was diluted with methylene chloride (120 ml) and washed with 0.5 M HCl solution (75 ml), water (50 ml), and brine (50 ml). The organic layer was separated, dried over sodium sulfate and concentrated under vacuum. The resulting residue was purified by column chromatography over silica gel (80 g) eluting with a methylene chloride/methanol gradient. Pure product containing fractions were combined and concentrated under vacuum to give 2',3'-di-*O*-benzyloxycarbonyl-5'-deoxy-4'-fluoro-5'-iodouridine (4.2 g, 94% yield) as a white solid.

$^1$H NMR (400 MHz, $CDCl_3$) δ 9.02 (s, 1H), 7.44 – 7.28 (m, 10H), 7.14 (d, *J* = 8.0 Hz, 1H), 5.86 – 5.72 (m, 2H), 5.69 – 5.57 (m, 2H), 5.19 (d, *J* = 4.3 Hz, 2H), 5.09 (d, *J* = 3.1 Hz, 2H), 3.71 – 3.35 (m, 2H).

$^{19}$F NMR (376 MHz, $CDCl_3$) δ -107.06 (td, *J* = 18.6, 7.3 Hz).

2',3'-Di-*O*-benzyloxycarbonyl-4'-fluorouridine (Fig. S17). In a 100 ml round-bottomed flask a 55% tetrabutylammonium hydroxide solution in water (8.04 ml, 9.37mmol) was adjusted to pH 3.5 by dropwise addition of trifluoroacetic acid (0.72 ml, 9.37 mmol) while maintaining a temperature below 25°C. The mixture was then treated with a methylene chloride (15 ml) solution of 2',3'-di-*O*-benzyloxycarbonyl-5'-deoxy-4'-fluoro-5'-iodouridine (2 g, 3.12mmol) followed by addition of 3-chloroperbenzoic acid (3.6 g, 15.62 mmol) in portions over a 30 minutes period. After 1 hour the pH drifted to pH 1.4. The mixture was adjusted back to pH 3.5 with 1N sodium hydroxide and allowed to stir for 16 hours after which time tlc (10% methanol in methylene chloride) and LCMS indicated complete conversion. The reaction mixture was quenched by addition of sodium thiosulfate (3.21 g, 20.31 mmol) slowly in portions while

maintaining a temperature below 25°C. After stirring for 30 minutes, the methylene chloride layer was separated, and the aqueous layer extracted with additional methylene chloride (2 × 30 ml). Combined organic layers were dried over sodium sulfate, concentrated, and purified by column chromatography over silica gel (80 g) eluting with 60% ethyl acetate in hexanes followed by a second column of silica gel (80 g) eluting with a methylene chloride/methanol gradient to give 2',3'-di-*O*-benzyloxycarbonyl-4'-fluorouridine (1.05 g , 63% yield) as a white solid.
$^1$H NMR (400 MHz, $CDCl_3$) δ 9.30 (s, 1H), 7.39 – 7.29 (m, 10H), 7.21 (d, *J* = 8.1 Hz, 1H), 5.83 (dd, *J* = 17.8, 7.0 Hz, 1H), 5.77 – 5.71 (m, 2H), 5.61 (dd, *J* = 7.0, 2.4 Hz, 1H), 5.17 (d, *J* = 4.8 Hz, 2H), 5.09 (s, 2H), 3.86 (q, *J* = 5.8, 4.9 Hz, 2H), 3.06 (s, 1H).
$^{19}$F NMR (376 MHz, $CDCl_3$) δ -121.03 (dt, *J* = 17.7, 4.6 Hz).

4'-Fluorouridine 5'-*O*-triphosphate (Fig. S17). A 10 ml round-bottomed flask charged with 2',3'-di-*O*-benzyloxycarbonyl-4'-fluorouridine (348 mg, 0.66 mmol) and anhydrous trimethyl phosphate (3.5 ml). After stirring for 20 minutes at room temperature, the solution was cooled to 0°C and treated with 1-methyl-imidazole (115 µL, 1.44 mmol) followed by dropwise addition of phosphorus oxychloride (122 µL, 1.31 mmol) over a 40-minute period. The mixture continued to stir at 0°C for 3.5 hours after which time tlc (10% methanol in DCM and then 7:2:1 *i*PrOH:$NH_4OH$:water) indicated complete phosphorylation. The mixture was treated with tributylamine (0.94 ml, 3.94 mmol), tris(tetrabutylammonium)pyrophosphate (887 mg, 0.98 mmol), and anhydrous DMF (1.5 ml). After 1 hour at RT, the reaction mixture was quenched with 100 mM TEAB (20 ml), stirred for 1 hour, degassed by pump-fill with argon (3x) and treated with 10% palladium on carbon (100 mg). After cooling with an ice-bath, the mixture was pump-filled with hydrogen (2×) followed by vigorous stirring under atm pressure of hydrogen for 30 minutes. After vacuum evacuation of the reaction vessel, the mixture was filtered through a pad of Celite and the collected palladium washed with water (2 × 20 ml). Combined filtrates were washed with ether (4 × 60 ml) and then concentrated under vacuum at 25°C. After co-evaporation with water (2 × 25 ml), the crude triphosphate was purified by column chromatography over DEAE-Sephadex GE A-25 (10 mm × 130 mm) eluting with a buffer gradient from 100 mM (450 ml) to 500 mM TEAB (450 ml). Pure product containing fractions as determined by tlc (8:1:1 $NH_4OH$:iPrOH:water) were combined and concentrated under vacuum with the bath temperature set at 25°C. The solid was dissolved in methanol (1 ml) and treated with saturated solution of sodium perchlorate in acetone (10 ml). The resulting white precipitate was collected by centrifuge, washed with acetone (5 × 5 ml), dissolved in water (1 ml), and concentrated by lyophilization to yield 4'-fluoro-uridine 5'-*O*-triphosphate (3.14 mg, 0.81% yield) as the tetrasodium salt.
$^1$H NMR (400 MHz, $D_2O$) δ 7.77 (d, *J* = 8.0 Hz, 1H), 6.15 (d, *J* = 1.9 Hz, 1H), 5.91 (d, *J* = 8.1 Hz, 1H), 4.72 – 4.57 (m, 1H), 4.41 (d, *J* = 6.3 Hz, 1H), 4.30 (ddd, *J* = 10.2, 6.3, 3.0 Hz, 1H), 4.17 (dt, *J* = 10.8, 5.0 Hz, 1H). $^{31}$P NMR (162 MHz, $D_2O$) δ -7.81(d), -11.84 (d, *J* = 19.2 Hz), -22.23 (t). $^{19}$F NMR (376 MHz, $D_2O$) δ -121.09 (unresolved dt, *J* = 19.2 Hz). LCMS Calculated for $C_9H_{13}FN_2O_{15}P_3$ [M-H$^+$]: 500.9; found: 500.8

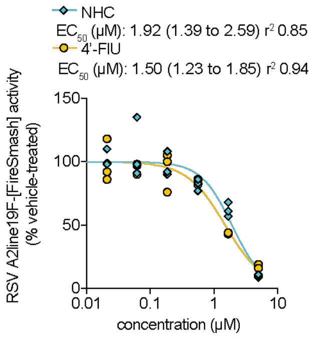


**Fig. S1.**

**Dose-response curves of 4'-FlU and NHC against recRSV-A2line19F-[FireSmash] on HEp-2 cells.** Viral inhibition determined by reporter activity. 4-parameter variable slope regression modeling, $EC_{50}$ values and 95% CIs with goodness of fit ($r^2$) are shown. Symbols represent independent repeats (n=3).

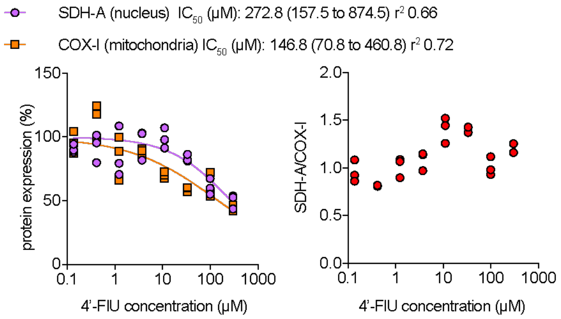


**Fig. S2.**

**Dose-dependent measure of 4'-FlU on mitochondrial biogenesis.** HBTEC cells were assayed for reduction in nuclear encoded protein SDH-A or mitochondrial-encoded protein COX-I after 48-hour incubation with 3-fold serial dilutions of 4'-FlU from 300 µM or vehicle (DMSO) treatment. 4-parameter variable slope regression modeling. $CC_{50}$ values and 95% CIs with goodness of fit ($r^2$) are shown. Symbols represent mean and error bars standard deviation (n=3).

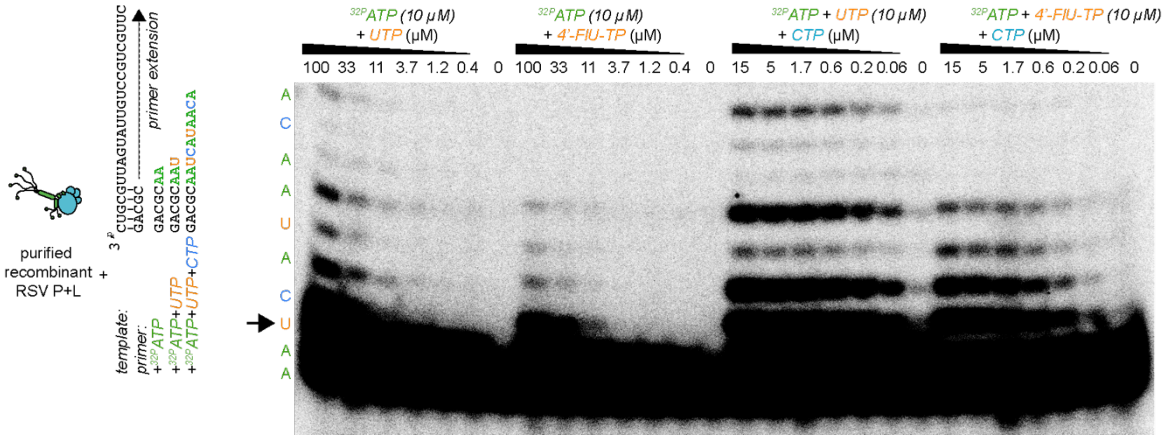


**Fig. S3.**

**Increased contrast of Fig. 2C.** Gel is annotated according to Fig. 2C.

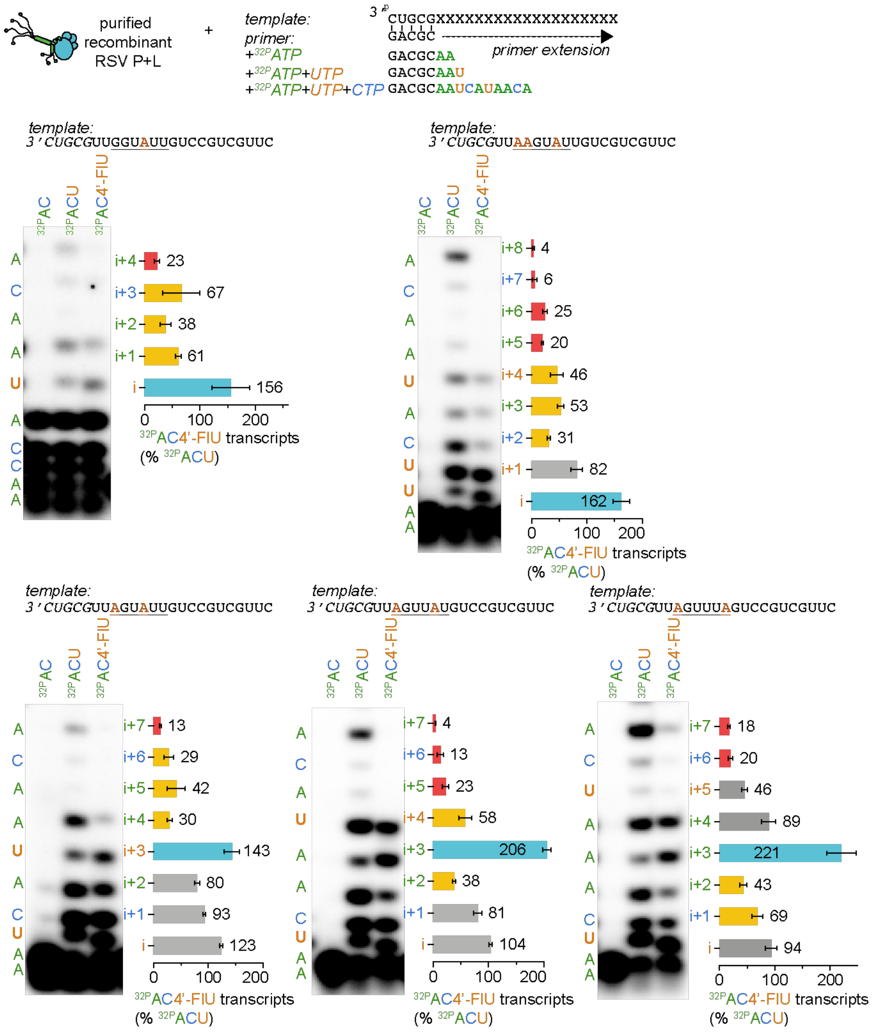


**Fig. S4.**

**Additional templates for RSV primer extension assay.** Gels are annotated according to Fig. 2.

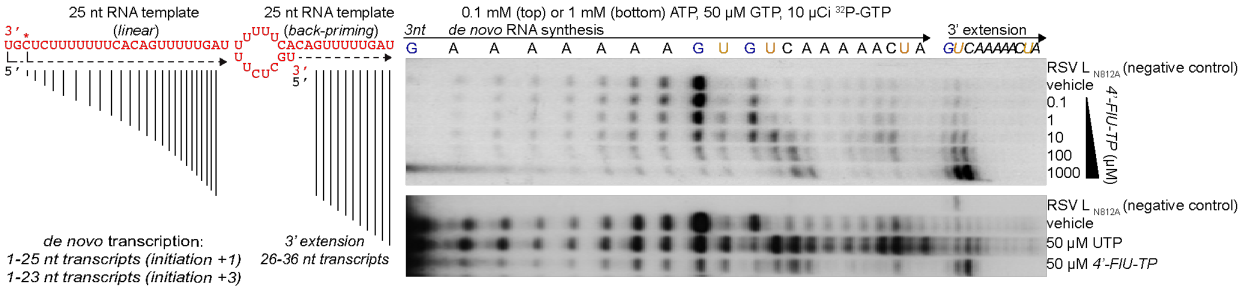


**Fig. S5.**

***In vitro* activity of 4'-FlU-TP on *de novo* RNA synthesis by RSV RdRP.** A 25-nucleotide synthetic RNA template corresponding to an authentic RSV promoter sequence (trailer complement) is sufficient to promote RNA synthesis by purified recombinant RSV RdRP complexes. RNA transcripts generated contain full 25-nucleotide transcripts with all intermediates resulting from aborted transcription products, and longer 26-36-nucleotide transcripts corresponding to a primer extension driven by stochastic back-priming of the template on itself. Reactions were made in presence of indicated concentration of nucleotides (n=1).

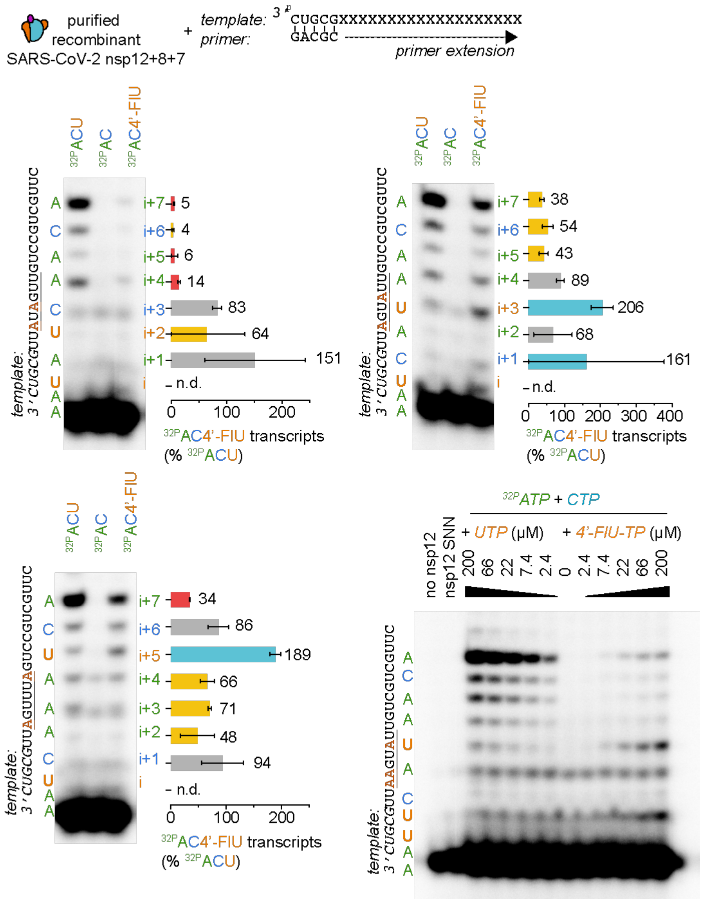


**Fig. S6.**

**Additional templates for SARS-CoV-2 primer extension assay.** Gels are annotated according to Fig. 2.

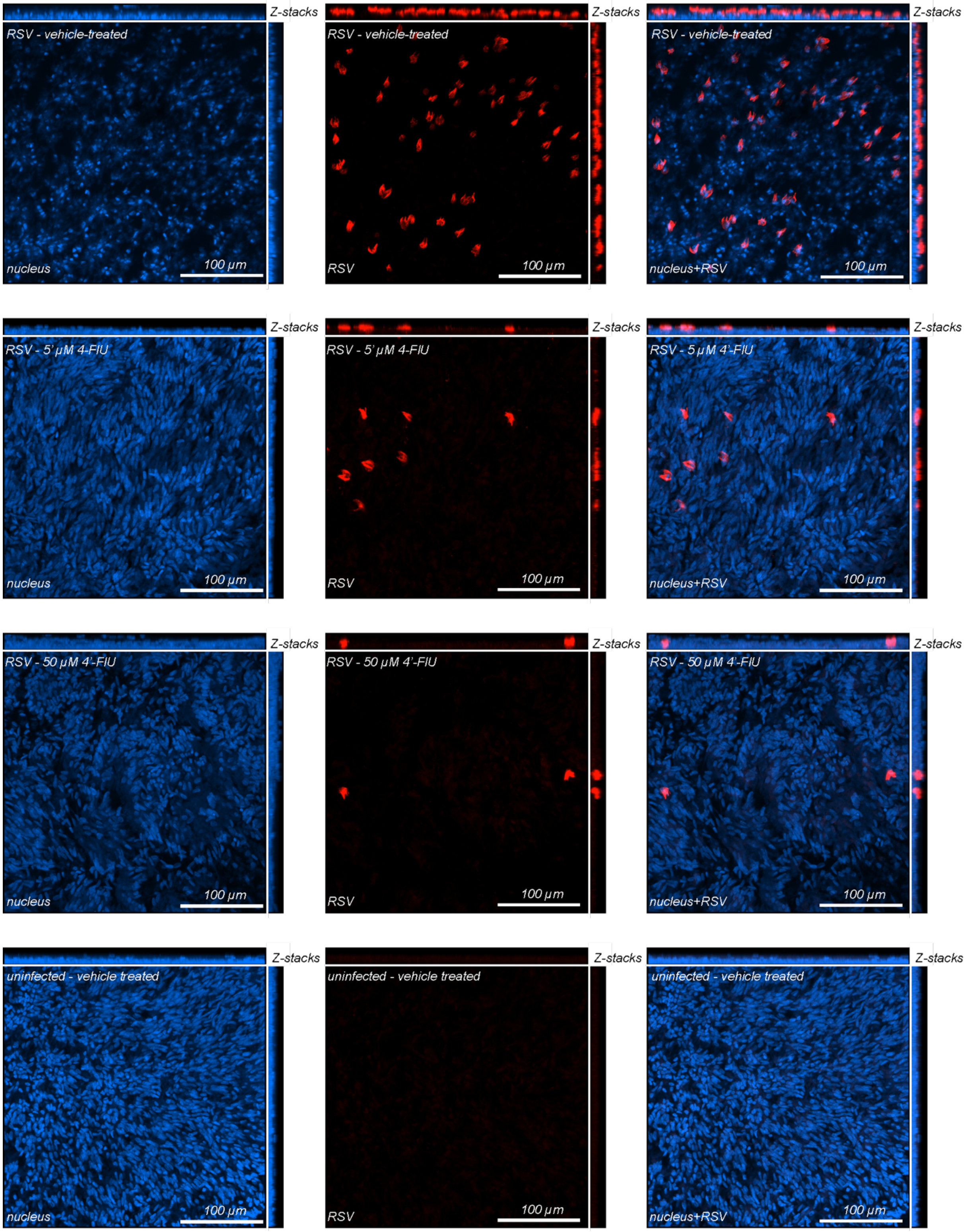
Z-stacks
RSV - vehicle-treated
nucleus
100 µm
Z-stacks
RSV - vehicle-treated
RSV
100 µm
Z-stacks
RSV - vehicle-treated
nucleus+RSV
100 µm
Z-stacks
RSV - 5' µM 4-FlU
nucleus
100 µm
Z-stacks
RSV - 5' µM 4-FlU
RSV
100 µm
Z-stacks
RSV - 5 µM 4'-FlU
nucleus+RSV
100 µm
Z-stacks
RSV - 50 µM 4'-FlU
nucleus
100 µm
Z-stacks
RSV - 50 µM 4'-FlU
RSV
100 µm
Z-stacks
RSV - 50 µM 4'-FlU
nucleus+RSV
100 µm
Z-stacks
uninfected - vehicle treated
nucleus
100 µm
Z-stacks
uninfected - vehicle treated
RSV
100 µm
Z-stacks
uninfected - vehicle treated
nucleus+RSV
100 µm

**Fig. S7.**

**4’-FlU visually decreases the presence of detectable RSV-positive cells in differentiated HAE.** Cells from Fig 3F were observed with a lower 20x objective to better appreciate the density of infected cells. At 5 µM and 50 µM treatment RSV-positive cells were sporadically detected. Representative pictures showing staining for RSV infected cells (red) and nucleus (blue). z-stacks of 20 µm slices (1 µm thick) with 20× objective. Dotted lines represent the location of x-z and y-z stacks. Scale bar: 100 µm.

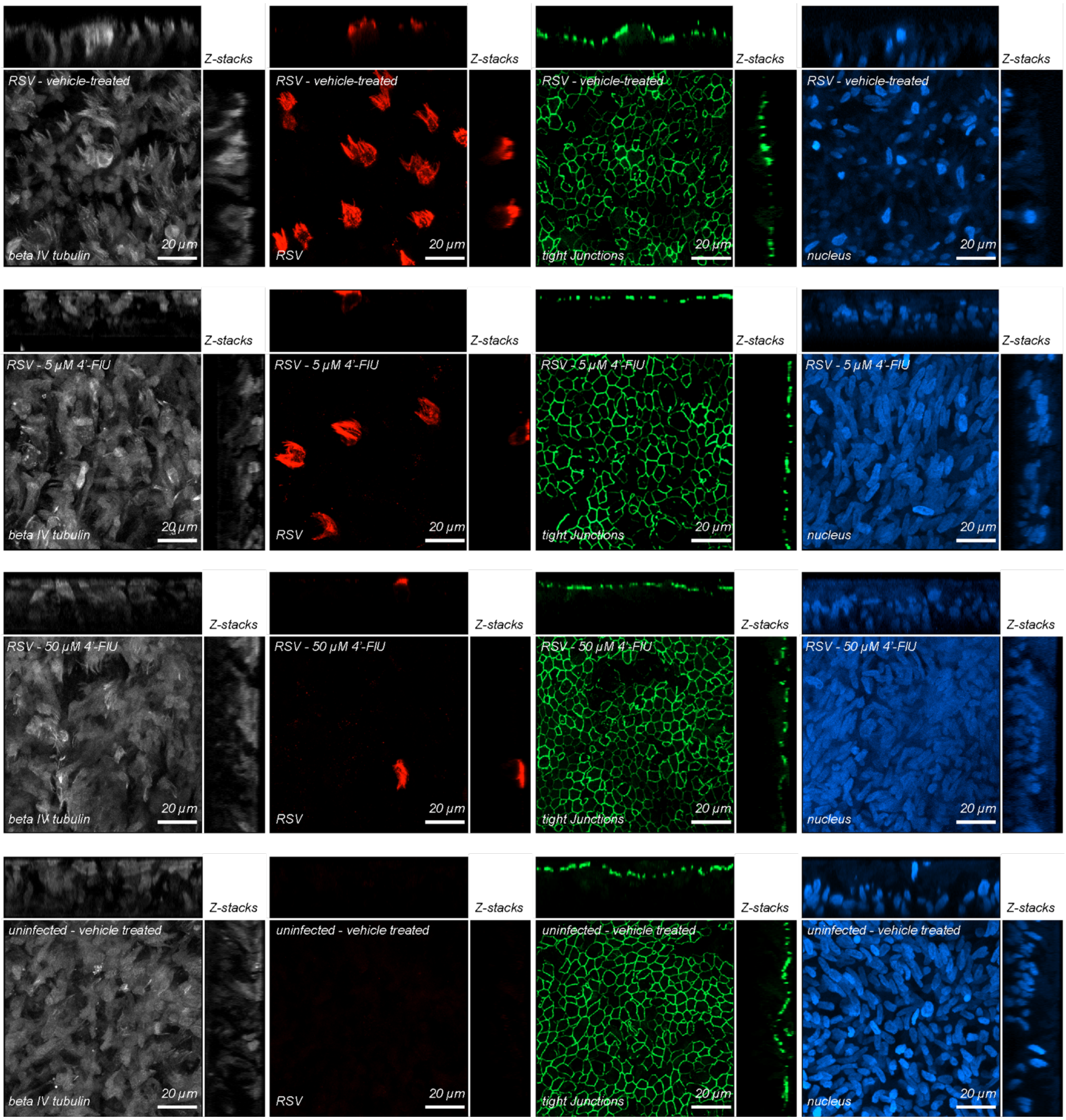


**Fig. S8.**

**4'-FlU does not alter the pseudostratified organization of differentiated epithelium.** Each channel of the overlays represented in Fig 3F-H are separated: RSV infected cells, tight junctions and nuclei were stained with anti-RSV immunostaining, anti-ZO-1 immunostaining and Hoechst

34580 and are colored in red, green and blue, respectively. Absent from the overlays, ciliated cells were stained through anti-beta tubulin IV immunostaining and are colored in white.

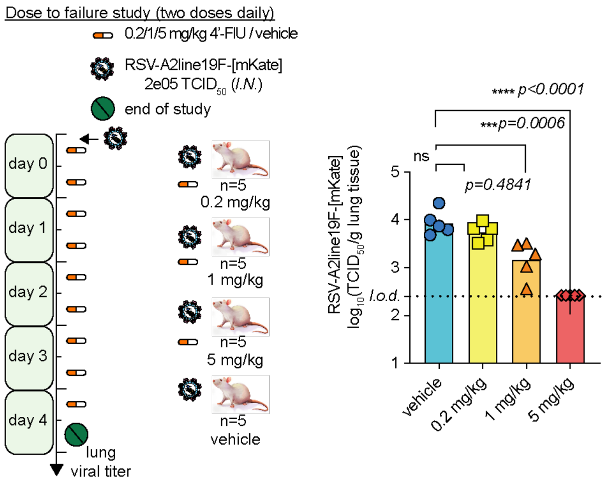


**Fig. S9.**

**Dose to failure study with treatment twice daily.** 6-8 weeks old female Balb/cJ mice were intranasally inoculated with 200,000 $TCID_{50}$ recRSV-A2line19F-[mKate] and treated orally with 0.2, 1 or 5 mg/kg mouse body weight of 4'-FlU (formulated in 10 mM sodium citrate and 0.5% Tween80 in water) or vehicle twice daily starting at 2 hours post-infection (A). At day 4.5 post-infection, viral lung titers were determined with $TCID_{50}$ titration on HEp-2 cells (B). Symbols represent individual values and bar represent means viral titers. One-way ordinary ANOVA with Dunnett's post-hoc multiple comparisons (n=5).

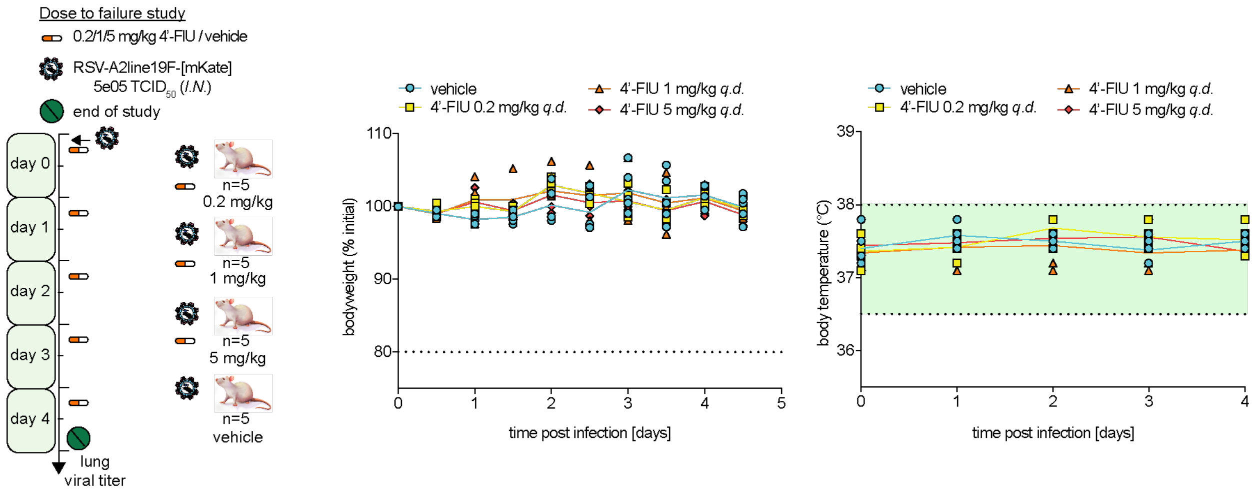


**Fig. S10.**

**Body weight and body temperature of mice from Fig. 4A-B.** Symbols represent individual values (n=5).

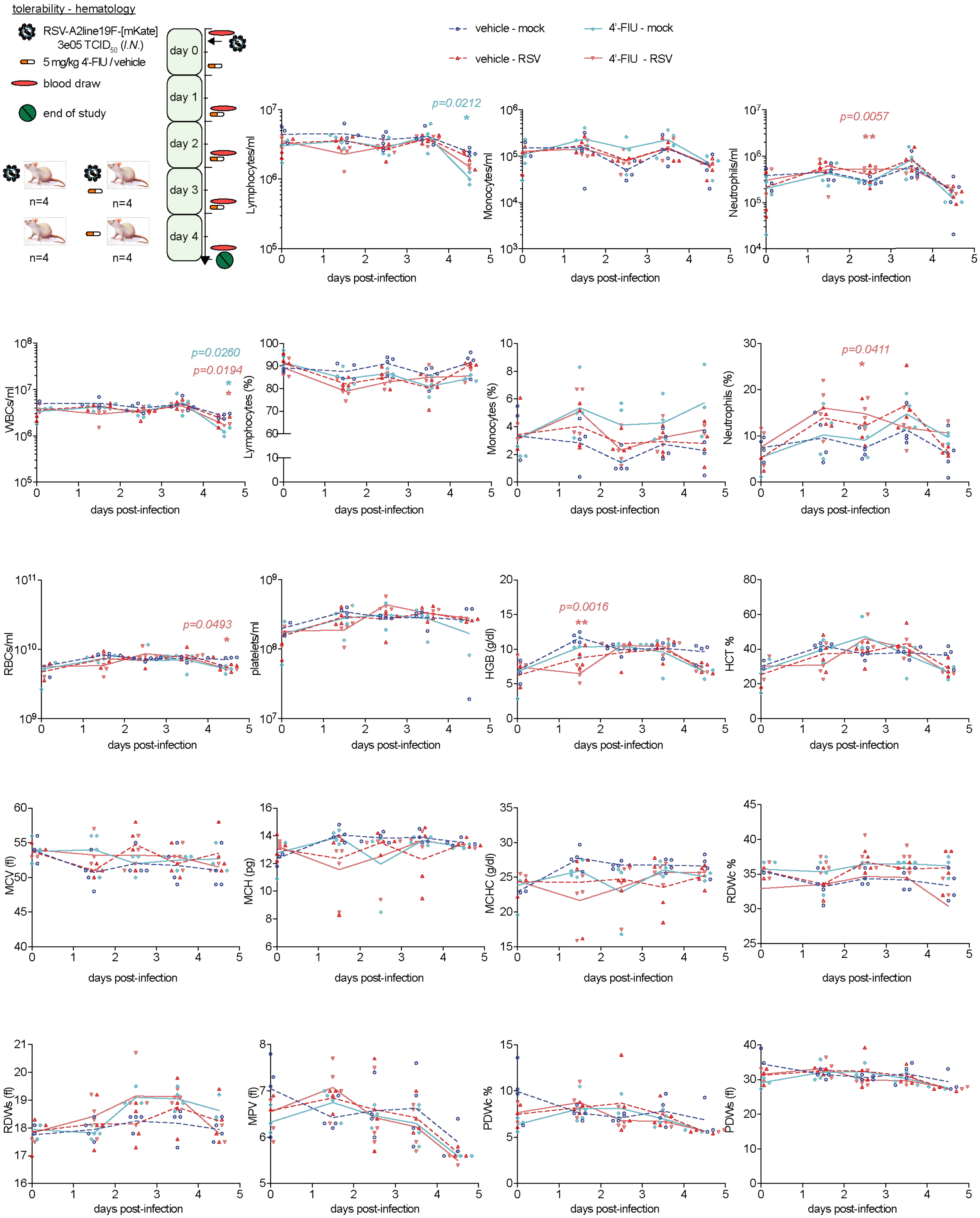
tolerability - hematology
RSV-A2line19F-[mKate]
3e05 TCID50 (I.N.)
5 mg/kg 4'-FIU / vehicle
blood draw
end of study
day 0
day 1
day 2
day 3
day 4
n=4
n=4
n=4
n=4
vehicle - mock
4'-FlU - mock
vehicle - RSV
4'-FlU - RSV
p=0.0212
p=0.0057
p=0.0260
p=0.0194
p=0.0411
p=0.0493
p=0.0016
Lymphocytes/ml
Monocytes/ml
Neutrophils/ml
WBCs/ml
Lymphocytes (%)
Monocytes (%)
Neutrophils (%)
RBCs/ml
platelets/ml
HGB (g/dl)
HCT %
MCV (fl)
MCH (pg)
MCHC (g/dl)
RDWc %
RDWs (fl)
MPV (fl)
PDWc %
PDWs (fl)
days post-infection

**Fig. S11.**

**Complete blood count parameters from Fig. 4C-D.** Symbols represent individual values (n=4).

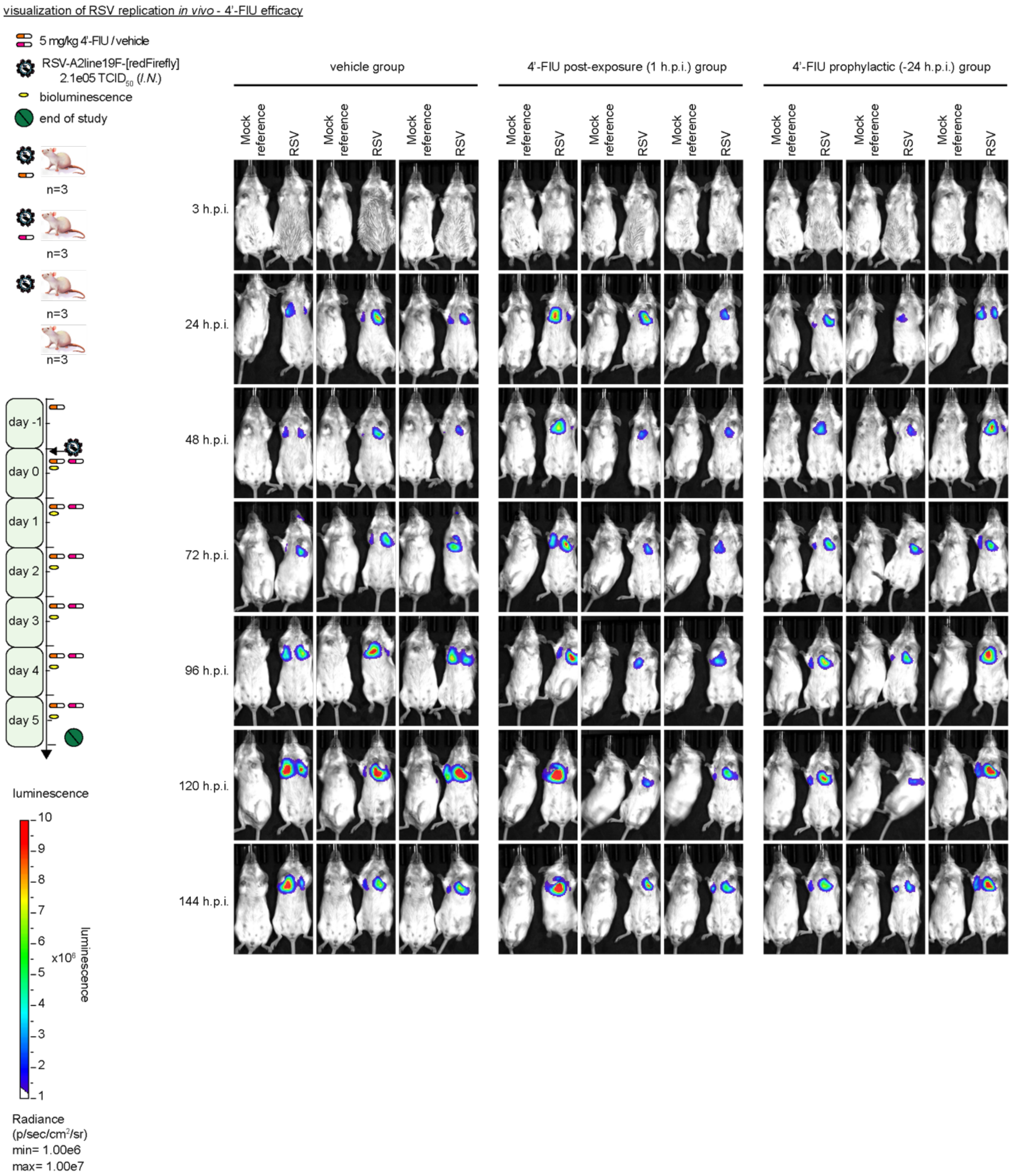


**Fig. S12.**

**Live imaging of 4'-FlU efficacy against RSV replication in mice lungs at days 0-5 post-infection.** 6-8–week old female Balb/cJ mice were intranasally inoculated with 210,000 $TCID_{50}$ recRSV-A2line19F-[redFirefly] and treated orally with 5 mg/kg mouse body weight of 4'-FlU (formulated in 10 mM sodium citrate and 0.5% Tween80 in water) once daily starting at either

24 hours prior to infection or 1 hour post-infection. Starting at 3 hours post-infection, luciferase activity was measured with an IVIS system (Caliper) under isoflurane anesthesia. 100 µl of 100 mg/ml D-luciferin (Millipore-Sigma) was retro-orbitally injected after administration of a single drop of propacaine and signal measured with a sequence of 9×30 seconds starting at 30 seconds post-substrate injection.

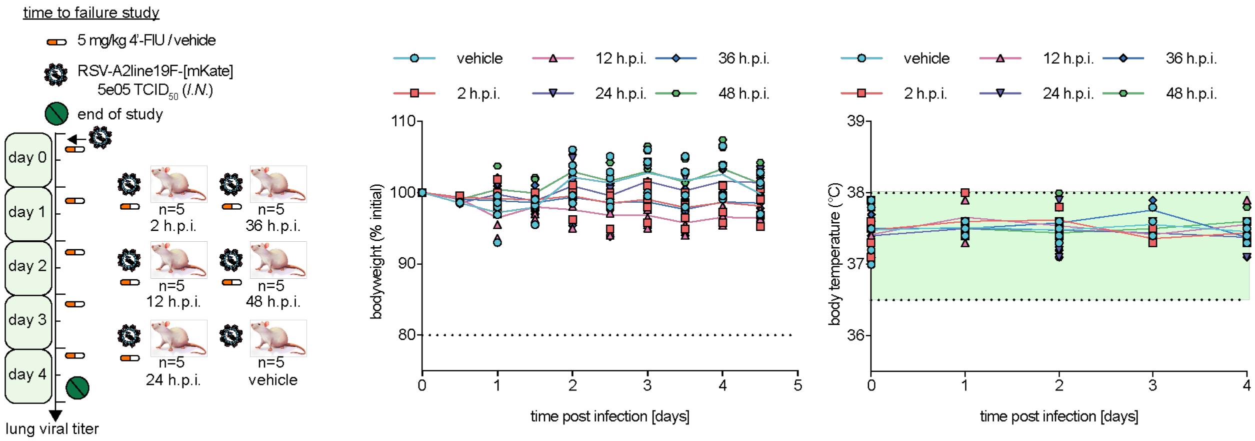


**Fig. S13.**

**Body weight and body temperature of mice from Fig. 4A-B.** Symbols represent individual values (n=5).

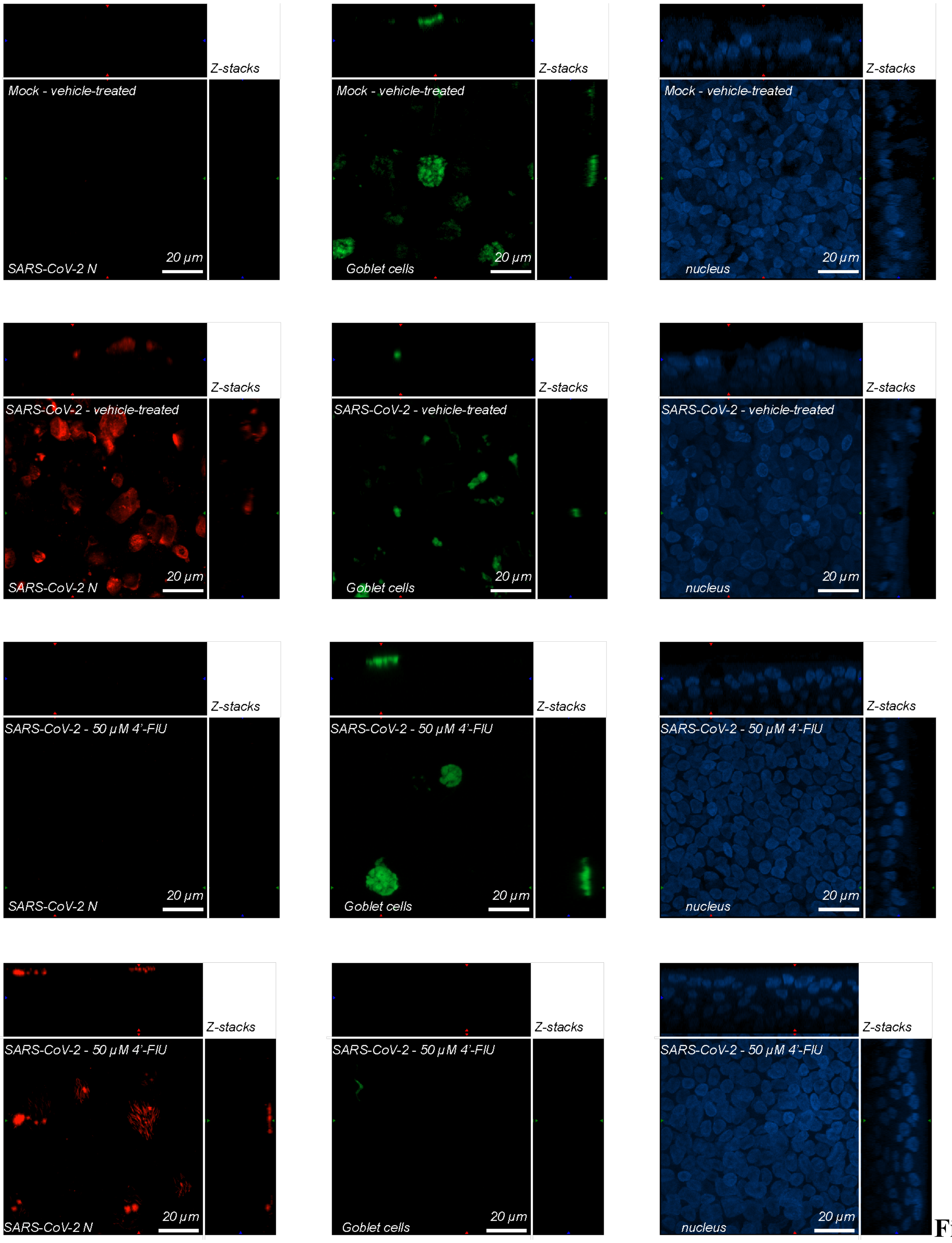


**Fig. S14.**
**4'-FlU does not alter the pseudostratified organization of SARS-CoV-2 infected differentiated epithelium.** Each channel of the overlays represented in Fig. 5B,C,E,F are

separated: SARS-CoV-2 infected cells, Goblet cells and nuclei were stained with anti-SARS-CoV-2 N immunostaining, anti-MUC5AC immunostaining and Hoechst 34580 and are colored in red, green and blue, respectively.

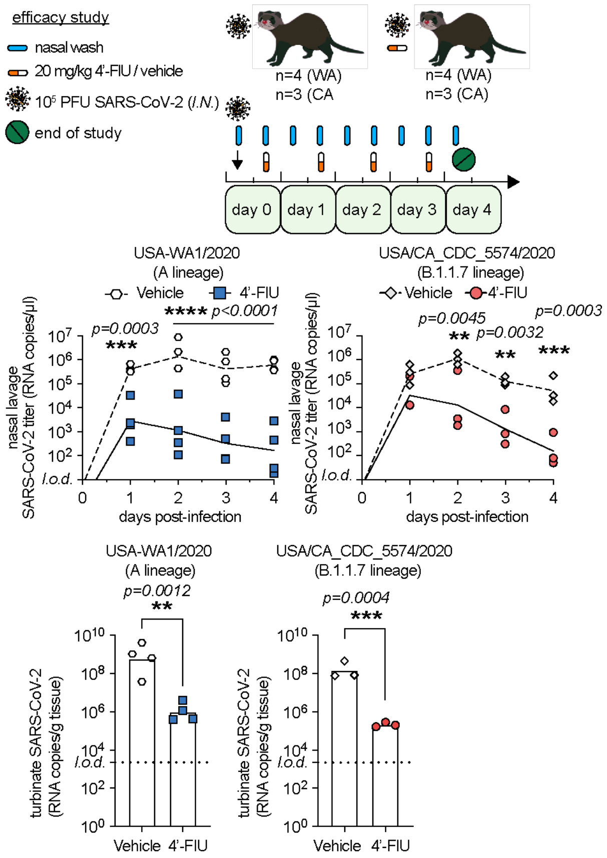


**Fig. S15.**

**SARS-CoV-2 RNA copies number in nasal washes and turbinate samples from Fig.6.** Annotated as in Fig. 6.

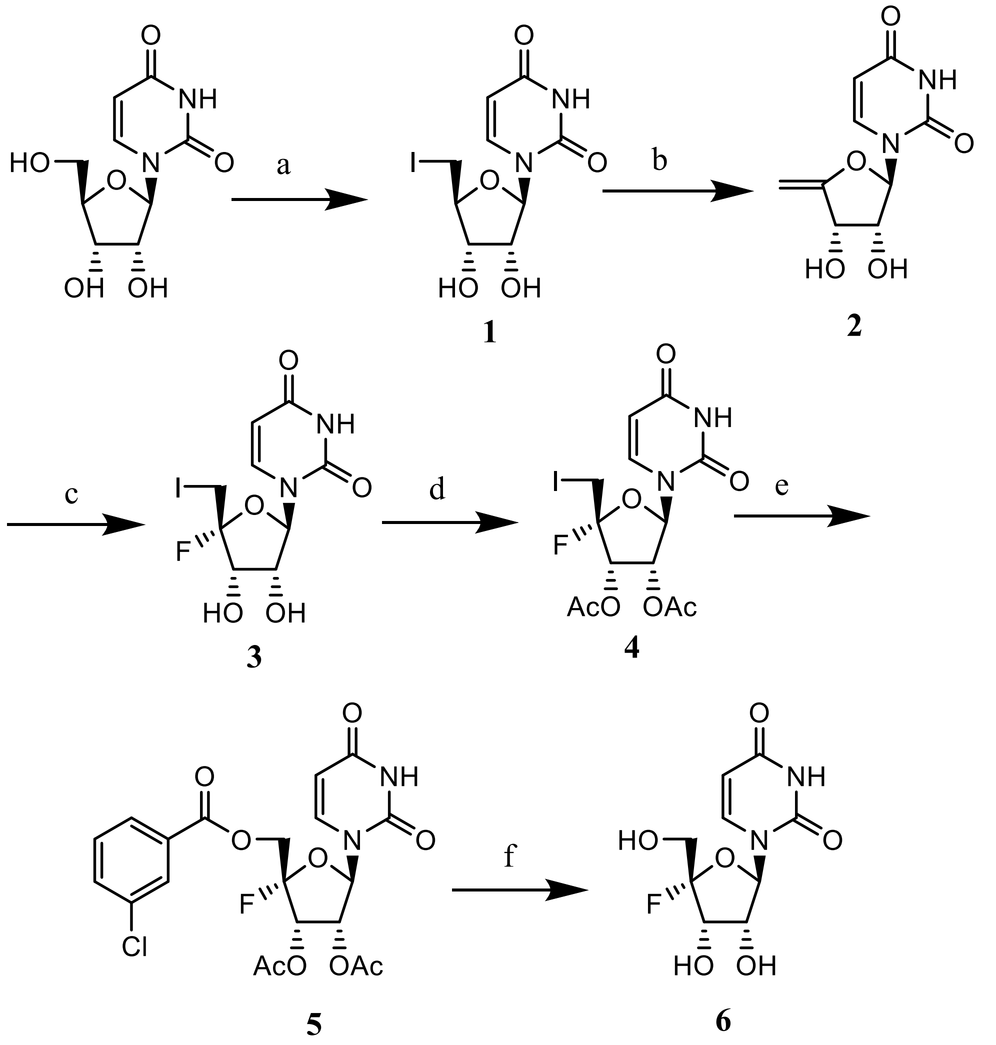


**Fig. S16.**

**Chemical synthesis.** 5’-deoxy-5’-iodouridine synthesis scheme. a) I2, $PPh_3$, Imidazole, THF; b) NaOMe, MeOH; c) $Et_3N$-3HF, NIS, $CH_3CN$; d) $Ac_2O$, $Et_3N$, DMAP, $CH_2Cl_2$; f) $Bu_4NHSO_4$, 3-Chlorobenzoic acid, MCPBA, $K_2PO_4$, water, $CH_2Cl_2$; f) 4N $NH_3$, $CH_3OH$

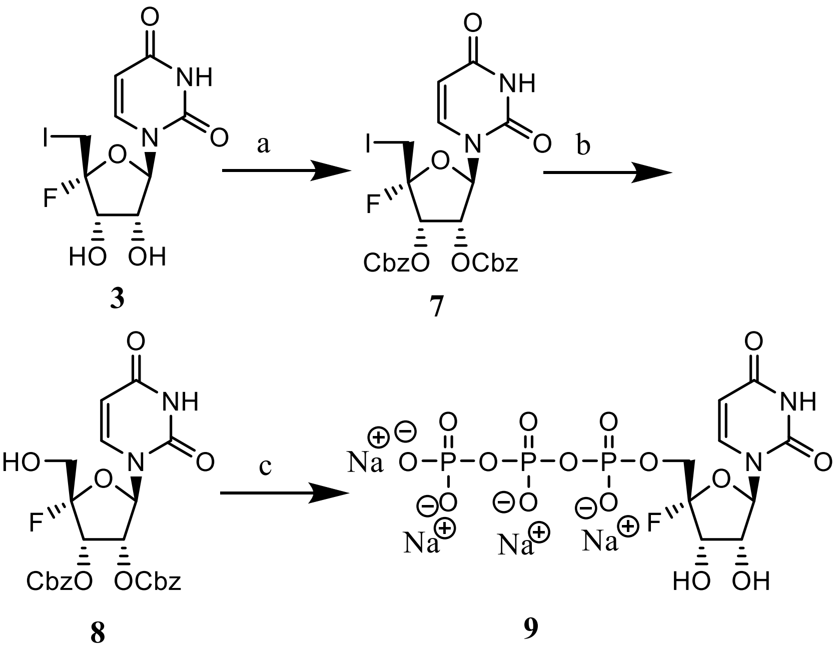


**Fig. S17.**

**Chemical synthesis.** a) CbzCl, 1-Methylimidazole, $CH_2Cl_2$; b) 55% aqueous $Bu_4NOH$, TFA, MCPBA, $CH_2Cl_2$; c) 1. $P(O)Cl_3$, 1-Methylimidazole, $(O)P(OCH_3)_3$; 2. $Bu_3N$, $(Bu_4N)_{33}+P_2O_{73}$-H, DMF; 3. $H_2$, 10% Pd/C

| **Viral target or minireplicon source** | **$EC_{50}$ [µM]** | **95% CI [µM]** | **Cell line or type** | **Assay readout** |
|---|---|---|---|---|
| RSV isolate 6A8 (B) | 0.61 | (-) | HEp-2 | Virus titration |
| RSV isolate 16F10 (B) | 0.67 | (-) | HEp-2 | Virus titration |
| RSV isolate 2-20 (A) | 0.66 | (-) | HEp-2 | Virus titration |
| recRSV-A2line19F-[mKate] | 1.20 | (-) | HEp-2 | Virus titration |
| recRSV-A2line19F-[FireSmash] | 0.07 | 0.06-0.09 | HAE ("F1") | Luciferase reporter |
| recRSV-A2line19F-[FireSmash] | 0.09 | 0.05-0.14 | HAE ("M4") | Luciferase reporter |
| recMeV-[$N_{PEST}$luc] | 0.10 | 0.09-0.6 | Vero/hSLAM | Luciferase reporter |
| recHPIV-3-[Nluc] | 0.41 | 0.39-0.45 | VeroE6 | Luciferase reporter |
| recSeV-[Gluc] | 0.04 | 0.03-0.05 | VeroE6 | Luciferase reporter |
| recVSV-[Nluc] | 0.64 | 0.6-0.68 | VeroE6 | Luciferase reporter |
| recRabV-[Nluc] | 0.16 | 0.14-0.18 | VeroE6 | Luciferase reporter |
| SARS-CoV-2 USA-WA1/2020 | 5.1 | (-) | VeroE6 | Virus titration |
| SARS-CoV-2 USA/CA_CDC_5574/2020 | 0.46 | (-) | VeroE6 | Virus titration |
| RSV minireplicon | 3 | 1.96-4.57 | HEK-293T | Luciferase reporter |
| MeV minireplicon | 0.04 | 0.02-0.09 | BSR-T7/5 | Luciferase reporter |
| NiV minireplicon | 1.81 | 1.01-3 | BSR-T7/5 | Luciferase reporter |
| HPIV-3 minireplicon | 0.33 | 0.22-0.49 | BSR-T7/5 | Luciferase reporter |
| recRSV-A2line19F-[FireSmash] | 1.5 | 1.23-1.85 | HEp-2 | Luciferase reporter |
| recRSV-A2line19F-[FireSmash] | 0.055 | (-) | HAE ("F1") | Virus titration |
| SARS-CoV-2 | 2.47 | (-) | HAE ("F1") | Virus titration |

**Table S1**

**$EC_{50}$ concentrations of 4'-FlU against different target viruses.**

| Cell line or type | Cell culture media | $CC_{50}$ [µM] | 95% CI [µM] |
|---|---|---|---|
| MDCK | DMEM | >500 | (-) |
| HEp-2 | DMEM | >500 | (-) |
| BSR-T7/5 | DMEM | >500 | (-) |
| BEAS-2B | DMEM | >500 | (-) |
| BEAS-2B | RPMI | >500 | (-) |
| BEAS-2B | RPMI glucose-free + galactose | 250 | 183-348 |
| HAE (“F1”) | BronchiaLife | 467.9 | 360-954.3 |
| HAE (“M4”) | PneumaCult EX-plus | 169 | 92.39-396.3 |

**Table S2**
**$CC_{50}$ concentrations of 4’-FlU.**

| Ferret | Dose [mg/kg] | $t_{max}$ [h] | $C_{max}$ [nmol/ml] | AUC-INF [h*nmol/ml] | $C_{max}$/Dose [kg*nmol/ml/mmol] | AUC-INF/Dose [h*kg*nmol/ml/mmol] | $t_{1/2}$ [h] |
|---|---|---|---|---|---|---|---|
| **1** | **15** | 1 | 28.6 | 144.5 | 499.3 | 2525.4 | 5.6 |
| **2** | **15** | 1 | 34.2 | 132.3 | 598.7 | 2312.2 | 6.3 |
| **3** | **15** | 1 | 41.6 | 185.1 | 726.7 | 3236 | 6 |
| **mean** | **15** | **1.0 ± 0.0** | **34.8 ± 6.5** | **154.0 ± 27.6** | **608.2 ± 114.0** | **2691.2 ± 483.7** | **5.97 ± 0.35** |
| **4** | **50** | 1 | 68.3 | 452.1 | 358 | 2370.7 | 4.2 |
| **5** | **50** | 4 | 50.7 | 464.1 | 266 | 2433.4 | 3.5 |
| **6** | **50** | 2 | 70.9 | 323.1 | 372 | 1694.3 | 5.1 |
| **mean** | **50** | **2.3 ± 1.5** | **63.3 ± 10.9** | **413.1 ± 78.1** | **332.0 ± 57.5** | **2166.1 ± 409.8** | **4.3 ± 0.80** |

**Table S3.**

**Selected single oral dose PK properties of 4'-FlU in ferrets.** Samples were analyzed by a qualified LC-MS/MS method, calculations with WinNonlin; means ± SD (n=3).

**Data S1. (separate file)**

PDF document containing all uncropped gels (SDS-PAGE and RNA PAGE) and their biological repeats used in this study, annotated. Dotted lines show inserts used in the labeled figures.

**Data S2. (separate file)**

Worksheet containing all raw numerical data and statistical analysis of this study.
